# Cellular Reprogramming of H3K27M Pediatric High-Grade Glioma to Neuron-like State

**DOI:** 10.1101/2025.08.05.668496

**Authors:** Abicumaran Uthamacumaran, Cynthia Horth, Eric Bareke, Michel Gravel, Jacek Majewski

## Abstract

This study explores the cell fate reprogrammability of H3K27M-mutant pediatric high-grade gliomas (pHGG) using neuronal transdifferentiation as a potential targeted therapy. We treated the BT245 patient-derived glioma cell line with pharmacological combinations targeting neuronal differentiation pathways and performed bulk RNA sequencing to characterize gene expression patterns driving cell fate transitions. Our findings reveal that the drug combinations induce transcriptomic changes consistent with differentiation towards neuronal phenotypes, including the upregulation of synaptic and dendritic signaling genes and the downregulation of malignant signatures. In comparison, astrocytic differentiation media (DM) and H3K27M knockout (KO) promote residual astrocytic phenotypes, suggesting neuronal transdifferentiation as a more effective strategy for mitigating tumor aggressiveness and progression. Differentially expressed genes such as GRIK1, GRIN1, NRXN3, NRXN1, CALB2, SCGN, SLC32A1, SLC1A2, KCNC3, and neurodevelopmental regulators including WNT7A, DLX6, ERBB4, ARX, BCL11B, SEMA3C, and FGFBP3 were identified as key markers regulating the neuron-like lineage transition. This study demonstrates that pHGGs can be phenotypically redirected toward neuronal-like identities through modulating cell fate differentiation programs. These findings advance the concept of ‘differentiation therapy’ as a promising intervention to reduce phenotypic plasticity and malignancy in pHGG ecosystems. While these are early *in vitro* findings, the ability to steer and control glioma cells toward stable, less malignant fates offers promising translational potential for patient-centered targeted therapies.

## Introduction

H3.3K27M-mutant Diffuse Midline Gliomas (DMGs) are an aggressive molecular subtype of pediatric high-grade gliomas (pHGGs) and represent one of the leading causes of cancer-related death in children [1, 2]. Despite significant advances in understanding of the disease etiology through the identification of validated molecular drivers, multiomics, and data science, patient outcomes remain dismal, with a median survival of less than two years [1, 3]. The poor prognosis of pHGGs can be attributed to their nature as complex adaptive systems, characterized by emergent behaviors such as phenotypic plasticity, therapeutic resistance, invasiveness, and intratumoral heterogeneity [4–7].

Among the key driver mutations in DMG is the H3.3K27M oncohistone, which profoundly alters brain developmental trajectories by disrupting lineage-specific differentiation programs [3, 8–9]. The K27M mutation in histone H3 variants, such as the H3.3 in the BT245 cell line, impairs trimethylation at lysine 27 (H3K27me3) by inhibiting the activity of the Polycomb Repressive Complex 2 (PRC2). This drives a global reduction in repressive epigenetic marks across key developmental genes, contributing to dysregulated neurodevelopment [3, 8, 10].

Single-cell transcriptomic studies further reveal that pHGGs navigate a hybrid continuum of transcriptional states, co-expressing distinct lineage markers in ways that confer phenotypic plasticity [5, 7, 11]. The resulting developmental blockade traps cells in critical tipping points, i.e., unstable stem-like intermediate states, disrupting neurodevelopmental hierarchies [12, 13]. This instability drives maladaptive behaviors, such as heterogeneity, invasion and therapy resistance, which are key features of HGGs [14, 15]. Earlier studies suggest that H3.3K27M-mutant DMGs arise from developmentally arrested or “stalled” differentiation of oligodendrocyte precursor cells (OPCs), leading to epigenetically trapped progenitor states [3, 14, 16]. Meanwhile, another epigenetically distinct pHGG subtype, the H3.3 G34R/V gliomas, develops from GSX2/DLX-expressing interneuron progenitors [17]. However, recent single-cell analyses by Wang et al. (2025) revealed that H3.3K27M DMGs also share transcriptomic features with intermediate neuronal progenitor cells (IPCs), suggesting a more fluid neurodevelopmental spectrum [15]. Thereby, we hypothesized that although DMGs predominantly arise in the brainstem, they may retain a developmental (lineage) plasticity that could be harnessed to promote neuron-like reprogramming.

Challenging the notion of a strict lineage-specific plasticity in HGG systems, ‘differentiation therapy’ or ‘cellular reprogramming’ is an emerging strategy in precision oncology that attempts to overcome cancer plasticity by redirecting tumor cells toward more stable, specialized, and less proliferative lineages [18], or cancer reversion toward benign-like states [12, 19–21]. In the context of pHGGs, which are marked by developmental arrest and ‘stuck’ in stem-like or progenitor-like instability, promoting terminal differentiation, especially into non-dividing neuron-like cells, may reduce malignancy and invasiveness [18]. For example, pharmacological combinations such as the FIIDC (FID for short) and FTT cocktails have been shown to reprogram adult glioma cell lines toward neuronal-like lineage differentiation [22, 23]. More recently, exogenous Neurexin-1 (NRXN1) has been shown to induce neuron-like differentiation in adult glioblastoma stem-like cell lines (GSCs), suppressing mesenchymal traits and improving prognosis in mouse xenograft models [24].

Based on these findings, the present pilot study investigates the phenotypic reprogramming potential of BT245 glioma cells, a preclinical model for H3.3K27M mutant DMG. This study aims to identify lineage-specific programs and molecular biomarkers steering neural cell fate commitments in pHGGs by examining transcriptional changes induced by neuronal trans-differentiation drug combinations compared to astrocytic differentiation media (DM).

Furthermore, the effects of H3K27M knockout (KO) on these reprogramming pathways are analyzed to identify transcriptomic signatures associated with developmental blockades in cellular reprogramming. While the primary focus of this study is on neuronal differentiation as a reprogramming strategy, DM and KO conditions analyses serve as comparative experiments to contextualize the specificity and efficacy of the proposed neuronal approach. These conditions allow us to assess whether reprogramming toward an astrocytic lineage or removing the driver mutation alone can similarly suppress malignant traits. By comparing transcriptional trajectories across these conditions, we show that neuronal differentiation leads to more stable, and less invasive, whereas DM and KO result in either partial lineage shifts or retention of plastic states. We assessed the transcriptional effects of these reprogramming strategies through gene expression analysis to characterize lineage-specific transitions and phenotypic outcomes.

Together, our findings demonstrate that neuronal reprogramming can redirect pHGG cells toward lineage-committed states while constraining malignant plasticity and stem-like traits, highlighting its promise as a targeted therapy to suppress tumorigenicity in pHGG ecosystems.

## Materials and Methods

### Cell culture

BT245 tumor-derived cell lines from patients were cultured in Neurocult NS-A proliferation media (StemCell Technologies) with bFGF (10 ng/mL), rhEGF (20 ng/mL), and heparin (0.0002%) (all from StemCell Technologies). The plates were coated with poly-L-ornithine (0.01%) (Sigma) and laminin (0.01 mg/mL) (Sigma).

### Cellular Reprogramming

Two drug treatments (FID combo and FTT combo) were applied to the BT245 glioma cell line to induce neuronal differentiation. Sample collection timepoints were chosen based on phenotypic observations of cellular response. FID-treated cells showed early neuronal features around Days 3–4 and were harvested between Days 6–7. FTT-treated cells responded more slowly, with morphological changes evident closer to Day 10. Controls were collected at Day 5 to match the midpoint of induction schedules. These timepoints were selected empirically by visual observations of changes in microscopy, given this is the first study to test these differentiation cocktails in BT245 cells.

### Neuronal Trans-differentiation Drug Cocktails

The details of the sample preparation and cell culture methods are provided by Lee et al. (2018) and Gao et al. (2019) [22, 23]. In brief, there are two drug treatments: FID and FTT combinations.

### FID Treatment

The FID cocktail consisted of five small molecules: DAPT (5 µM, MedChemExpress), Forskolin (10 µM, MedChemExpress), IBET151 (2 µM, MedChemExpress), Laduviglusib/CHIR99021 (3 µM, MedChemExpress), and ISX9 (10 µM, MedChemExpress). Each compound was dissolved in DMSO to prepare 10 mM stock solutions and stored at −20°C. Prior to use, stocks were thawed and briefly warmed to 37°C to ensure complete dissolution [22]. FID treatments were applied in a 2:1 ratio of BrainPhys™ Neuronal Medium (StemCell Technologies) to NeuroCult™ proliferation media. An equivalent concentration of DMSO, matching that used in the drug-treated conditions, was added to vehicle control groups to account for solvent-related effects. See Table 1 for cocktail composition and final concentrations and Table 2 for treatment timeline and media conditions.

**Table 1.** Summary of drug components, and dosages for FID and FTT treatment cocktails used for neuronal reprogramming in BT245 glioma cells.

| **FIIDC (FID)**<br>**Treatment** | **Effects** | **Dosage** |
| --- | --- | --- |
| Laduviglusib<br>(CHIR99021) | GSK 3a/b inhibitor | 3 uM |
| ISX9 | Calcium channel activator for neural induction | 10 uM |
| Forskolin | Upregulates adenylate cyclase/cAMP agonist | 10 uM |
| DAPT | γ-secretase inhibitor | 5 uM |
| IBET151 | BET bromodomain inhibitor | 2 uM |
| **FTT**<br>**Treatment** | **Effects** | **Dosage** |
| Fasudil | Rho Kinase inhibitor | 50 uM |
| Tranilast | TGF-beta inhibitor | 100 uM |
| Temozolomide | Alkylating agent; Chemotherapy | 50 uM |

**Table 2.** Summary of treatment conditions, media compositions, and sample collection timepoints for BT245 glioma cells. This table outlines each experimental group, including drug treatments (FID or FTT), corresponding media formulations (BrainPhys and NeuroCult ratios), and timing of RNA sample collection as shown in Figure 1 schematic. Media adjustments was made for FTT Late sample to support viability at later timepoints in pediatric glioma cells, as increased cell death was observed. [<u>Note:</u> The parentheses under Sample Names denote the names as they appear in the GEO Repository (see Data Availability section)].

| Sample Name | Treatment Type | Drugs Used | Media Composition | Day of Collection |  |
| --- | --- | --- | --- | --- | --- |
| **BrainPhys**<br>(Ctr_Brain_1 / 2) | Control | DMSO only | 1/3 BrainPhys + 2/3 NeuroCult | Day 5 | Control (replicates) |
| **Neurocult**<br>(Ctr_Neuro_2) | Control | DMSO only | 100% NeuroCult | Day 5 | Control |
| **FTT Early**<br>(FTT_1) | FTT-treated | Fasudil, Tranilast, TMZ | 1/3 BrainPhys + 2/3 NeuroCult | Day 5 | Early FTT treatment |
| **FTT Late**<br>(FTT_Neuro_2) | FTT-treated | Fasudil, Tranilast, TMZ | 100% NeuroCult (switched in at Day 7) | Day 10 | Neurocult media added to support viability before RNA collection |
| **FID Early**<br>(FID_D6) | FID-treated | DAPT, Forskolin, IBET151, Laduviglusib, ISX9 | 2/3 BrainPhys + 1/3 NeuroCult | Day 6 | Captures early transcriptional shifts |
| **FID Late**<br>(FID1 / FID2) | FID-treated | DAPT, Forskolin, IBET151, Laduviglusib, ISX9 | 2/3 BrainPhys + 1/3 NeuroCult | Day 7 | Biological replicates |

### FTT Treatment

The FTT cocktail included Fasudil (50 µM, Selleckchem), Tranilast (100 µM, TCI Chemicals), and Temozolomide (50 µM, Selleckchem). Fasudil and Tranilast were dissolved in DMSO to prepare 173.87 mM stock solutions, and TMZ was prepared at 103.01 mM [23]. All stocks were gently heated to 50°C in a water bath and vortexed in microcentrifuge tubes to ensure dissolution, then aliquoted and stored at −20°C. FTT treatments were applied in a 1:2 ratio of BrainPhys™ Neuronal Medium to NeuroCult™ proliferation media, in contrast to the 2:1 ratio used for FID. These adjustments were made because Gao et al. [23] used BrainPhys only during FTT induction in later phases of reprogramming glioblastoma cells, whereas Lee et al. [22] applied 100% BrainPhys throughout the FID protocol. These media adjustments were also further tailored to support the viability and growth of our BT245 cells. Control groups received DMSO vehicles with volumes equivalent to those used in the drug-treated samples. For details on composition, refer to **Table 1**.

### Controls and Media Usage

The three control samples included two replicates cultured in BrainPhys:NeuroCult (1:2), denoted as BrainPhys, and one sample (denoted as Neurocult) maintained in 100% NeuroCult to represent an undifferentiated reference. All controls received matched DMSO volumes to mirror treatment groups. The differential use of BrainPhys (to promote neuronal differentiation) and NeuroCult (to preserve stem-like properties or viability) was based on protocols adapted from Lee et al. [22] for FID and Gao et al. [23] for FTT. Detailed timing, treatment conditions, and sample descriptions are summarized in Table 2.

### Time-Lapse Bright-Field Microscopy Imaging

Live-cell bright-field images were captured using the Bio-Rad ZOE™ Fluorescent Cell Imager at 20× magnification. BT245 glioma cultures were imaged at multiple time points during neuronal trans-differentiation using FID and FTT drug treatments, as well as untreated control conditions. Imaging was conducted at different time points to monitor morphological changes and guide RNA-seq sampling. Cells were imaged directly on culture plates without fixation or staining.

### RNA Sequencing and Bioinformatic Analysis

Total RNA was extracted and prepared for sequencing using the NEBNext® Ultra™ II Directional RNA Library Prep Kit (New England Biolabs), incorporating a ribosomal RNA (rRNA) depletion step to remove abundant rRNA species and enrich for both protein-coding messenger RNAs (mRNAs) and long non-coding RNAs (lncRNAs). This approach was selected to achieve more comprehensive transcriptome coverage than poly(A) selection-based methods. Library construction and sequencing were carried out at the McGill Genome Centre (Montreal, Canada), following the manufacturer’s protocols.

Bioinformatic processing of the RNA-seq data was conducted using the **bcbio-nextgen analysis pipeline** [https://bcbio-nextgen.readthedocs.io], which integrates a suite of best-practice tools for quality control, alignment, quantification, and statistical analysis. Initial quality assessment and trimming of raw reads were performed using **Atropos** [25], which removes sequencing adapters and low-quality bases. Quality control of both raw and trimmed reads was assessed using **FastQC** [https://www.bioinformatics.babraham.ac.uk/projects/fastqc/], **samtools** stats [26], and **Qualimap** [27]. Summarized quality control reports were generated using **MultiQC** [28], providing a unified overview of quality metrics across all samples.

High-quality reads were aligned to the human reference genome (UCSC hg38 build) using the **STAR** aligner [29], a splice-aware algorithm optimized for accurate and efficient mapping of RNA-seq data. Genome annotation was performed using a GTF file based on the standard **GENCODE** annotations (v38 – Release 38), used to guide transcript discovery and feature assignment.

Following alignment, gene-level quantification was performed using **featureCounts** [30], which assigns mapped reads to annotated genomic features (genes) in a strand-specific manner. The resulting count matrix was used as input for differential gene expression analysis.

Differential expression analysis was performed in R using the DESeq2 package [31], a widely used bioinformatics tool that models count data using negative binomial distribution. Genes with low read counts (< 10) across all samples were filtered prior to normalization and statistical testing. The dataset was normalized using DESeq2’s median-of-ratios method to account for sequencing depth and RNA composition. For each gene, log2 fold changes and statistical significance were estimated using the Wald test. Differentially expressed genes were defined as those with an absolute log2 fold change ≥ 2 and a Benjamini-Hochberg adjusted p-value < 0.05. The results were visualized and interpreted using a combination of volcano plots, hierarchical clustering, and principal component analysis (PCA).

For the neuronal differentiation protocol, bulk RNA-seq analysis detected 55,877 transcripts— including non-coding and predicted genes—across all samples. However, only a subset with sufficient expression (e.g., raw counts >10) was retained for downstream analysis. Using stringent criteria (|log2FoldChange| > 2 and adjusted p-value < 0.05 with Benjamini-Hochberg correction), we identified to 350 differentially expressed genes (DEGs) from the controls, of which 197 were upregulated and 153 were downregulated (Figure 3B).Applying such stringent cutoffs, particularly in small-sample RNA-seq datasets, helps prioritize substantially relevant expression changes while minimizing noise-driven false positives and enhancing specificity [32].

### Gene Set Enrichment Analysis and Cell Type Classification

For functional gene set enrichment, we used g: Profiler, an integrated web-based tool that supports data from Gene Ontology (GO), KEGG, Reactome, and other biological databases for using the list of identified DEGs [33]. g: Profiler performs statistical enrichment using the cumulative hypergeometric test and corrects for multiple testing using the Set Counts and Sizes (SCS) algorithm, which infers a dependency structure between functional categories. Ordered queries were enabled to allow enrichment analysis based on ranked gene lists (by log-fold change and smallest adjusted p-values), providing an additional layer of biological relevance [33].

In addition, we applied Fast Gene Set Enrichment Analysis (FGSEA) to identify biologically enriched pathways from gene expression data, using the C5: Gene Ontology (GO) Biological Processes gene set collection from MSigDB. FGSEA was run on two inputs: (1) the entire raw counts gene expression profile, and (2) the raw counts of DEGs identified by DESeq2 using the Wald test (padj < 0.05). For both analyses, genes were ranked by log2 fold-change. Fgsea was implemented using the fgsea R package with the following default parameters: minSize = 10, maxSize = 500, and nperm = 10,000. P-values were estimated using an adaptive multi-level split Monte Carlo scheme and adjusted for multiple testing using the Benjamini-Hochberg method. Fgsea has been benchmarked across over 600 datasets and shown to recover significantly enriched pathways with robust interpretation of transcriptomic data [34].

To infer cell-type identities from the DEG profiles, we used the Spatio-Temporal cell Atlas of the Brain, version 2 (STAB2), a database built from over 3.8 million single-cell and single-nucleus RNA sequencing profiles from the human and mouse brain [35]. STAB2 integrates developmental and anatomical annotations across 15 developmental stages and 63 brain subregions in humans (Yang et al., 2024), enabling predictive assignment of cell-type identities by comparing DEGs to known marker gene signatures derived from the curated transcriptomic clusters. Using hypergeometric test, STAB2 statistically evaluates the enrichment score of input gene sets (DEG list) against cell-type-specific markers, providing high-resolution annotations of cell types and their molecular subtypes. These enrichment scores were used to classify transcriptional states observed in our dataset [35].

### Network Inference and BDM Perturbation Analysis

To infer gene regulatory networks, DEGs from DSEQ2 were used to construct adjacency matrices based on Spearman correlations, with edges made absolute to avoid negative values. Network centrality measures (degree, betweenness, closeness, eigenvector, hub scores) were computed using the igraph package in R. Network visualization was performed with igraph and ggplot2, adjusting node sizes/colors to highlight key genes and scaling edge weights. The pyBDM package performed Block Decomposition Method (BDM)-based perturbation analysis, binarizing the network at a 0.5 threshold to identify causally linked genes impacting network robustness and resilience. BDM estimates the graph network’s algorithmic complexity to reveal markers driving neuronal lineage differentiation. Kolmogorov complexity predicts cell fate transitions by identifying network features and structural changes signaling critical phase transitions (e.g., bifurcations), and hence, provides a causal inference tool in network medicine [36].

## Results

### 1. Drug Treatments Promote Neuronal-like Structures in BT245 Cells

To investigate the neuronal differentiation potential in pediatric BT245 glioma cells, we treated the cell line with two distinct drug cocktails: FID (Laduviglusib, ISX9, Forskolin, DAPT, IBET151) and FTT (Fasudil, Tranilast, Temozolomide), previously demonstrated to induce neuronal phenotypes in adult high-grade glioma cell lines [22, 23] (see Methods). As shown in Figure 1, Day 1 marked the point of full confluence and the initiation of drug dosing.

**Figure 1.**
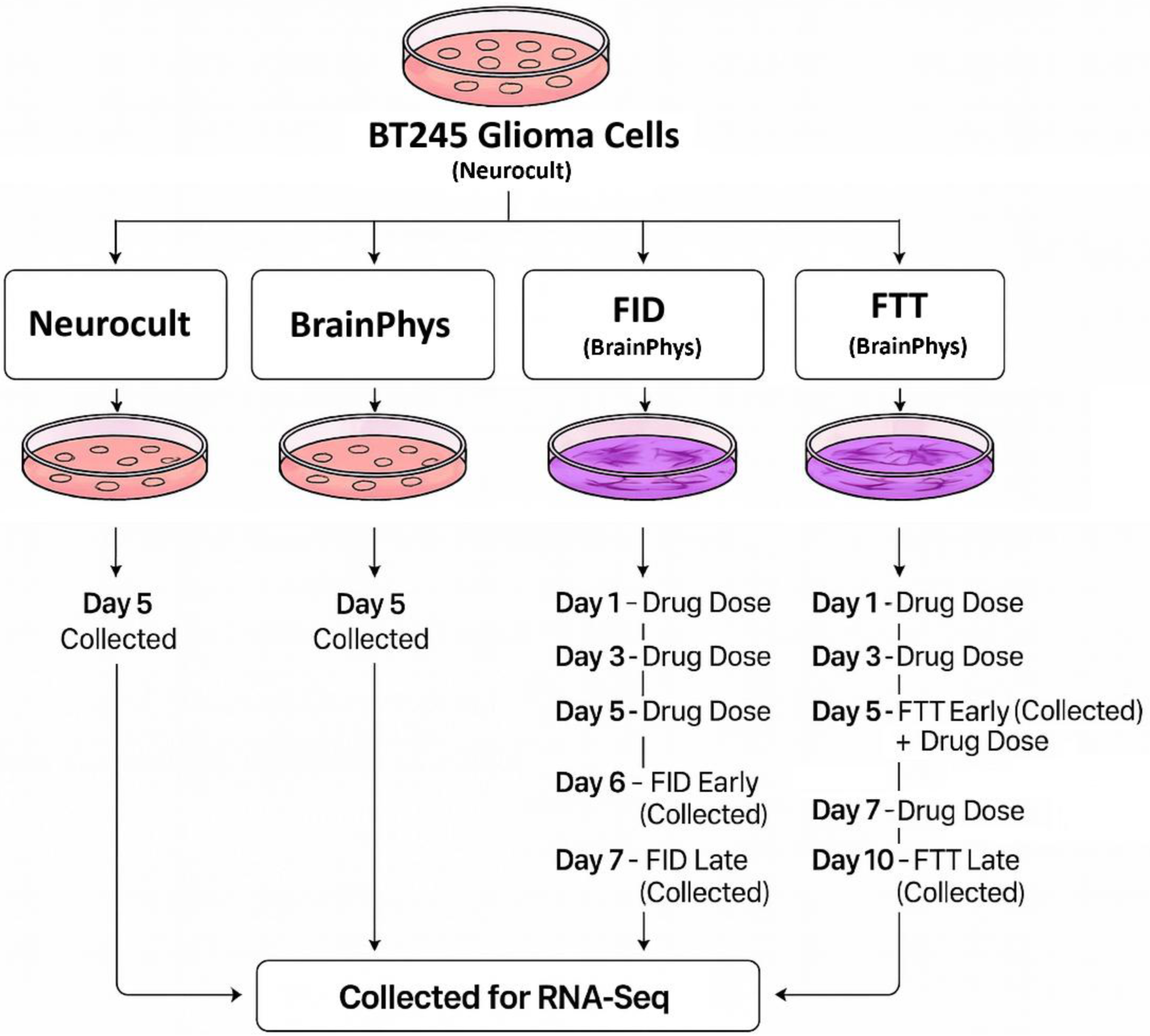
Schematic of Experimental Workflow. BT245 pediatric glioma cells were cultured under four conditions: two controls, namely 100% Neurocult (denoted as Neurocult) and a 1:2 ratio of BrainPhys:Neurocult (denoted as BrainPhys), and two differentiation treatments (FID and FTT). Day 1 corresponds to when cells reached full confluency and were exposed to the first drug dose. Media were refreshed with drug mixtures on Days 1, 3, and 5 for FID, and Days 1, 3, 5, and 7 for FTT groups. RNA-seq samples were collected at distinct time points to evaluate early and late responses: Day 6 (FID Early), Day 7 (FID Late), Day 5 (FTT Early), and Day 10 (FTT Late) for the treatment groups, and Day 5 for the control groups. <u>Note:</u> For FTT day 7 drug doses, we used 100% Neurocult instead of the 1:2 BrainPhys: Neurocult, to maintain cell viability.

Since the optimal drug treatment differentiation conditions required varying ratios of BrainPhys:Neurocult media, to determine the effects of media composition we cultured BT245 cells in two different control media conditions: NeuroCult and BrainPhys-based medium (1: 2 ratios of BrainPhys: NeuroCult). Neurocult represents standard maintenance media for BT245 cells, while the BrainPhys mixture follows the base media composition prescribed in the original differentiation protocols (treatments) [22, 23]. A 1:2 BrainPhys: Neurocult mix was specifically adapted for this cell line, as BT245 cells require Neurocult for viability. FID was dosed on Days 1, 3, and 5; FTT followed a similar schedule but exhibited delayed morphological responses. The drug-infused media were changed on these days (See Methods, Table 2 and Figure 1).

Bright-field imaging showed that both treatments induced neuron-like morphological changes in BT245 cells, while control cells retained undifferentiated morphology with no visible changes (Figure 2 A–B). FID treatment triggered early and pronounced differentiation: by Day 4 (Figure 2C), cells extended branched neuron-like projections, with changes stabilizing by Day 6–7 (Figure 2 D–E). Cell number and confluency were significantly reduced in FID-treated cells, indicating a marked decrease in surviving cells. By Day 7, FID-treated cells exhibited hallmark features of neuronal polarity and maturation, including thickened somas, long-range dendritic arborization, and fine axonal-like processes (Figure 2 E). In contrast, the FTT cocktail exhibited a delayed response, with mild neurite outgrowths only appearing around Day 7–10 (Figure 2 G-H). Occasionally cell death was observed, which may reflect treatment-associated stress. These morphological transitions support the hypothesis that FID, and to a lesser extent FTT, can reprogram glioma cells toward a stable, neuron-like fate.

**Figure 2.**
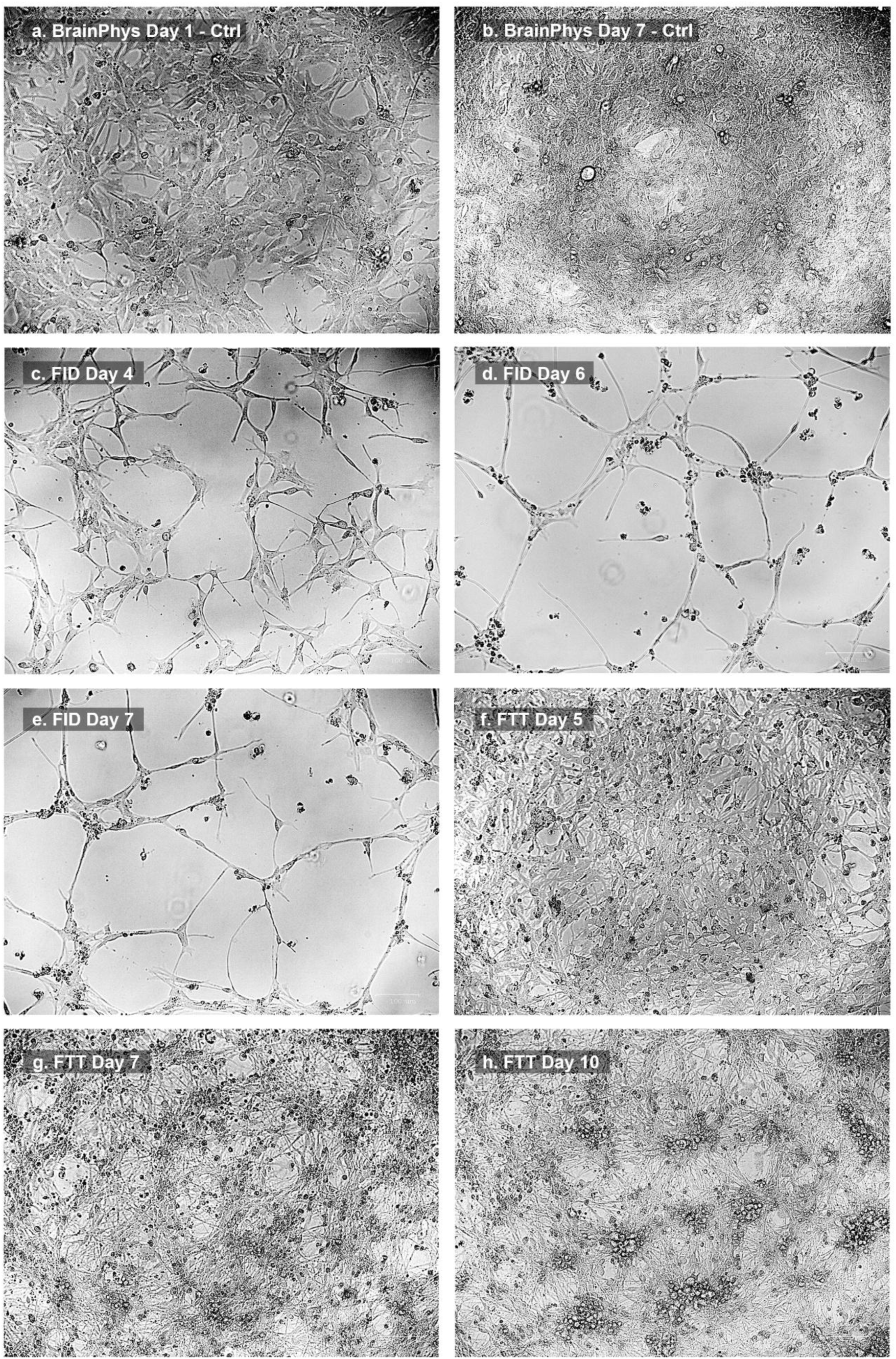
Microscopy Imaging of BT245 Controls and Drug Treatments. Control and FTT treated cells were grown in 1:2 ratio of BrainPhys:Neurocult media, and FID treated cells in 1:2 BrainPhys:Neurocult. Control cells (BrainPhys) at day 1 (A) and day 7 (B). No morphological changes from day 1 were observed in day 7. C) FID Treatment on day 4 showing some signs of neuronal-like differentiation. D) FID on day 6. FID-treated cells exhibited neuronal-like branching morphologies and were collected as the FID Early group. E) FID on day 7 (collected as FID Late group). F) FTT Treatment on day 5 (collected as FTT Early group). G) FTT on day 7. H) FTT on day 10 (collected as FTT Late for transcriptional profiling).

RNA-seq samples were collected at early and late timepoints post-treatment (Days 6 and 7 for FID; Day 5 and 10 for FTT, respectively), and on Day 5 for the controls. These time points were selected based on observed morphological changes under microscopy, as differentiation dynamics in BT245 cells differed from the original protocol timing, suggesting cell line specific changes. Throughout the treatment protocol, cells were not passaged; instead, only the drug-containing media was replaced at designated time points (Figure 1).

### 2. Neuronal Trans-Differentiation Treatments Reprogram BT245 Glioma Cells Toward a Neuronal-like Transcriptional Identity

Based on the observed morphological changes upon drug treatments, we selected appropriate time points as described above, to carry out transcriptomic analysis and characterize the differentiation processes at the molecular level. The effect of cellular reprogramming was evident in the global gene expression profiles. Principal component analysis (PCA) of the bulk transcriptome showed that control samples clustered together irrespective of the growth media used (Brainphys or Neurocult), while FID-treated and late-stage FTT-treated cells displayed distinct transcriptomic characteristics (Figure 3A). The FTT Early sample (Day 5) remained closer to the control cluster, consistent with its earlier differentiation stage observed during imaging (Figure 3A).

**Figure 3.**
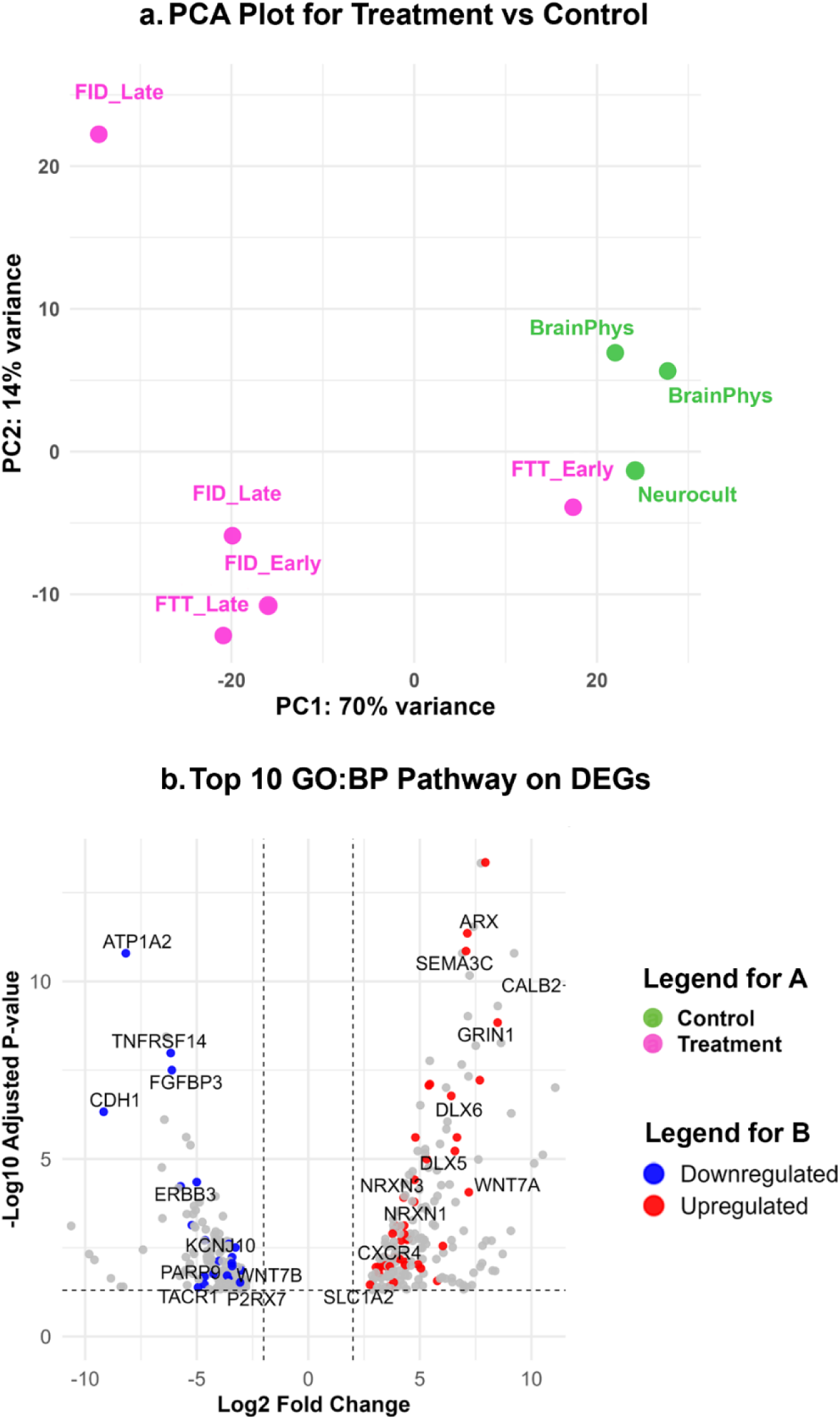
Effects of Neuronal Differentiation Drug Treatments on the BT245 Transcriptome. A) PCA clustering illustrates transcriptional shifts along the PC1 axis under FID and FTT treatments compared to controls. B) B) Volcano plot highlighting the top DEGs identified from fgsea (GO: BP) performed on DEGs only (padj < 0.05), using log2 fold-change ranking. These genes contribute to significantly enriched neuronal differentiation pathways.

To perform differential gene expression analysis, we compared the five combined FID and FTT treatment samples against the three control samples. Using stringent criteria (|log2FoldChange| > 2 and adjusted p-value < 0.05 with Benjamini-Hochberg correction), we identified to 350 differentially expressed genes (DEGs) from the controls, of which 197 were upregulated and 153 were downregulated (Figure 3B, Supplementary Table DEG). Applying such stringent cutoffs, particularly in small-sample RNA-seq datasets, helps prioritize substantially relevant expression changes while minimizing noise-driven false positives and enhancing specificity [32]. Visual inspection of the top DEGs identified many known markers of neuronal cell identity, such as GRIN1, DLX5, DLX6, ARX, NRXN1, NRXN3, SEMA3C, and CALB2.

To further contextualize these transcriptomic changes, we performed gene set enrichment analysis (fgsea) on the differentially expressed genes [34]. The top enriched pathways included key neurodevelopmental processes such as axon guidance, telencephalon development, and synaptic signaling (Table S1). Notably, among the top DEGs contributing to these pathways were established neuronal lineage markers and differentiation signatures such as GRIN1, NRXN1, DLX5, ERBB4, BCL11B, ARX, WNT7A, CALB2, SLC32A1, and KCNJ10 (Figure 3B).

### 3. Differentiation Treatments Reveal Neuronal Subtype-Specific Transcriptional Signatures and Reduced Expression of Malignancy-Associated Traits

Transcriptomic profiling revealed that differentiation treatments promote neurodevelopmental signatures encompassing both excitatory and inhibitory neuronal subtype identities (Figure 3B). To further support the neuronal differentiation observed in the transcriptomic data, we performed cell-type assignment analysis to infer the likely cellular identities and lineage transitions associated with the upregulated and downregulated DEGs. Using the Spatio-Temporal Atlas of the Brain (STAB-2), a curated reference of human and mouse brain cell types, we mapped DEGs to specific spatiotemporal lineages through marker-based annotation and hypergeometric enrichment testing [35] (see Methods). While we acknowledge that mapping bulk RNA-seq to single-cell-defined cell types may not fully capture transitional or hybrid (mixed) populations, this inference provides an approximation of lineage trajectories and dominant cellular identities.

STAB-2-based cell type classification [35] using upregulated DEGs suggests transcriptional similarity to excitatory neuron subtypes via the enrichment of excitatory neuron (ExN-TLE4) subtype markers in treated BT245 cells (Figure 4A). Among 197 upregulated genes, 73 map to ExN-TLE4 identity, including key signatures such as NRXN3, PTPRN, GRIN1, and CYP46A1. Inhibitory neuronal identity markers were also prominently upregulated. Among the 153 downregulated DEGs, many overlapped with glial lineages, such as astrocytic (e.g., AQP4), OPC (e.g., SOX10), and immune/mesenchymal (e.g., VEGFA, ILDR2, NR4A1) phenotypes (Figure 4B). These suppressed genes are associated with malignant traits such as migration, vasculature interaction, and extracellular matrix remodeling underlying invasion. Thus, the trend reveals a dual outcome of the treatments: a shift toward neuronal lineage identities, and concurrent silencing of glioma-promoting plasticity markers (reduced malignancy), highlighting the efficacy of the drug treatments in steering BT245 cells toward a stable, less tumorigenic phenotype.

**Figure 4.**
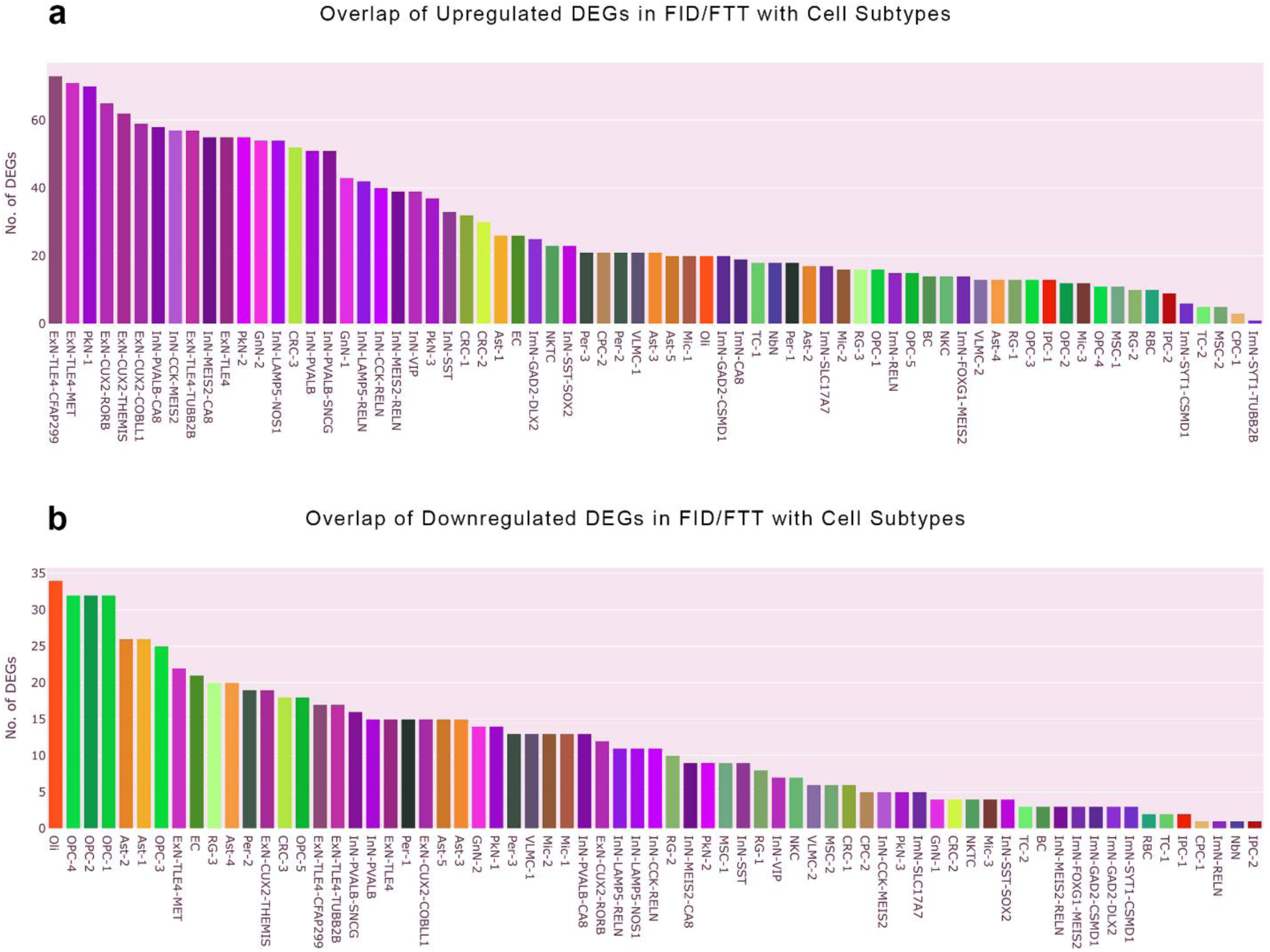
STAB-2 Cell Type Classification Reveals that Upregulated DEGs promote Neuronal phenotypes while Downregulated DEGs in Treatment Samples show a Reduction in Glioma malignancy. The bar plots show the number of DEGs matching signature genes of the listed cell types A) Upregulated DEGs, B) Downregulated DEGs. The assignments indicate that the treatment shifts the cellular identity towards excitatory neuronal types (A) and away from the oligodendrocytic/astrocytic lineages (B).

## 4. Neuronal Reprogramming Drives More Stable Lineage-Committed Transcriptional States than Astrocytic Differentiation and H3K27M KO

To assess the robustness and potential therapeutic advantages of neuronal differentiation (ND), we compared this strategy to two previously reported reprogramming approaches: astrocytic differentiation using serum-based differentiation media (DM) [37, 28] and H3.3K27M knockout (KO) in serum conditions [8] (See Supplementary Information). We used the RNA-seq data from those studies and compared the transcriptomic shifts across the ND, DM, and KO conditions. As shown in Figure 5A, ND-specific DEGs included neuronal identity markers consistent with neuronal lineage induction. In contrast, DM treatment led to strong upregulation of astrocytic and vascular genes such as GREM1, AQP4, and BMP5, while KO induced immune-modulatory and invasion-associated DEGs like TNFRSF11A, CD74, and CIITA.

**Figure 5.**
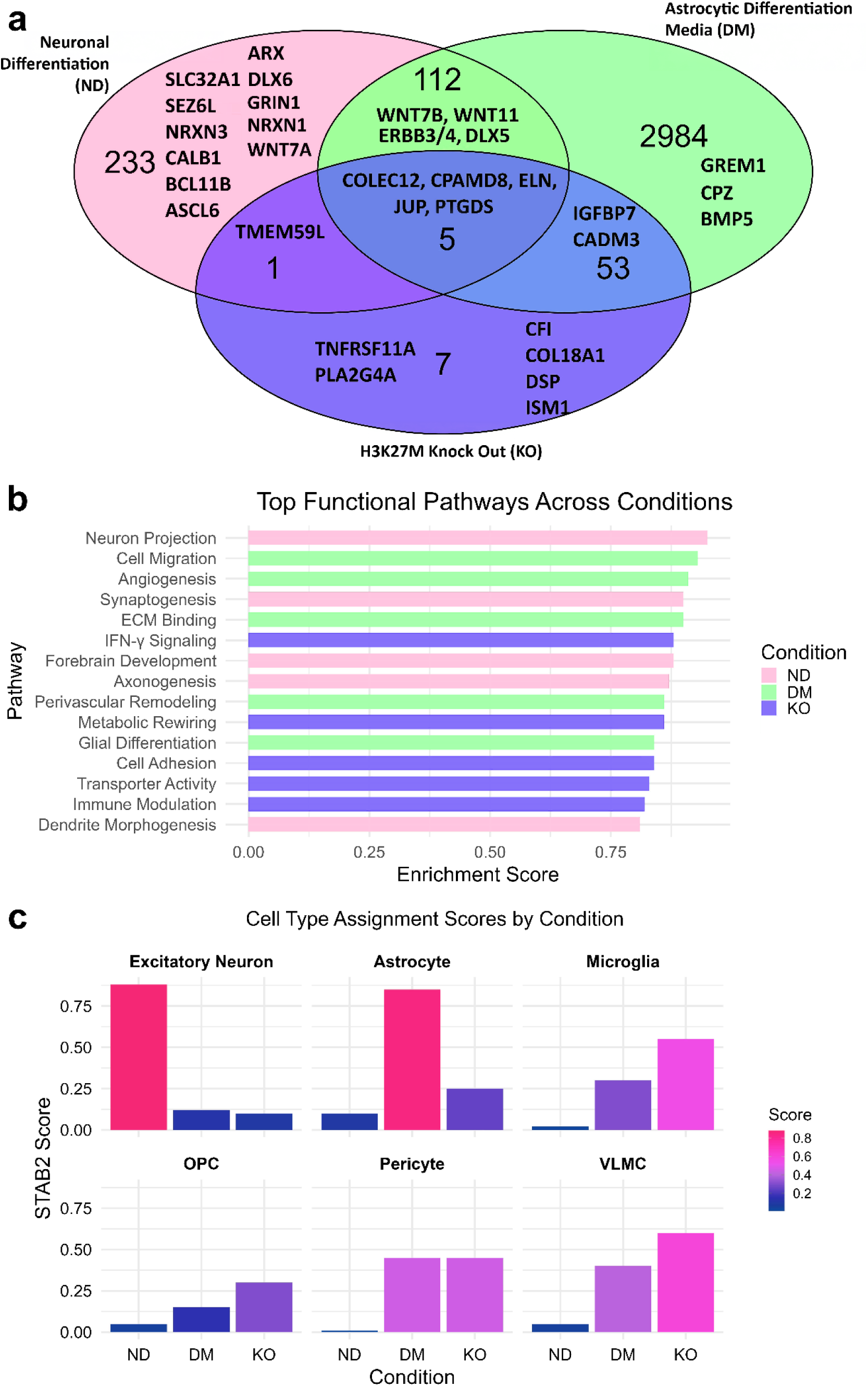
Comparative analysis of transcriptional programs induced by Neuronal Differentiation (ND), Astrocytic Differentiation Media (DM), and H3.3K27M Knockout (KO) in BT245 glioma cells. This figure summarizes the transcriptional and phenotypic reprogramming outcomes of three distinct interventions. ND refers to treatment with both neuronal differentiation cocktails aimed at reprogramming glioma cells toward neuronal lineages. DM involves exposure to serum-based astrocytic differentiation media to promote astroglial phenotypes. KO denotes CRISPR-mediated knockout of the H3K27M oncohistone to alleviate epigenetic barriers to differentiation and assess the resulting transcriptional changes. A) Venn diagram illustrating the overlap of significant DEGs among the three treatments. Key genes within each intersection are labeled. B) Bar plot showing the top Gene Ontology Biological Process pathways identified using fgsea applied to each condition’s log2FC-ranked DEGs. The enrichment score represents the normalized enrichment of a pathway within the ranked DEGs. C) STAB2-based cell type assignment scores computed from upregulated DEGs in each condition.

Fgsea-based pathway enrichment analysis (Figure 5B) further revealed divergent biological programs activated by each condition. As discussed above, ND strongly enriched neurodevelopmental pathways including neuron projection, axonogenesis, and synaptogenesis. DM activated signaling cascades involved in angiogenesis, extracellular matrix binding, and cell migration—hallmarks of a reactive astrocytic or vascular niche construction processes. KO treatment upregulated immune-inflammatory signaling, interferon gamma pathways, and metabolic rewiring, suggesting epigenetic de-repression of immune-related programs.

Lastly, STAB2-based cell type assignment analysis (Figure 5C) demonstrated that ND-treated cells most adopted an excitatory neuron-like identity, while DM-treated cells skewed toward astrocytic and pericyte-enriched phenotypes, and KO-induced programs reflected strong microglial and vascular-like (VLMC) identities. The STAB2 score represents a specificity-weighted enrichment based on reference single-cell datasets and highlights how the DEGs of each intervention drives transcriptional reprogramming toward distinct lineage fates.

Collectively, these findings demonstrate that while all three treatments modulate the glioma cell state, they navigate divergent lineage-specific mechanisms: ND promoting neuronal fate commitment, DM enhancing astrovascular characteristics, and KO activating immune-like and niche-adaptive responses. These findings suggest that ND potentially induces a less invasive, less proliferative profile consistent with neuronal lineage commitment. However, functional validation is needed to confirm maturation status, degree of plasticity, and whether the cell fate identity is subtype-specific or hybrid.

## Discussion

### 1. Evidence of Neuronal Features in Reprogrammed Glioma Cells using Neuronal Differentiation Cocktails

Our findings show that pharmacological induction of neuronal differentiation in H3K27M-mutant BT245 glioma cells resulted in robust upregulation of genes involved in neuronal projection, synaptogenesis, and radial glia-derived neuronal differentiation (Figure 3), supporting a transition toward a neuronal fate. Both FID- and FTT-treated cells exhibited transcriptomic signatures consistent with reduced malignancy and increased neuronal lineage commitment (Figure 4). Upregulated gene expression profiles were enriched for neuronal phenotypes associated with forebrain development and deep-layer cortical neurons. Further, cell type assignment using STAB2 revealed strong enrichment for neuronal subtypes involved in synaptic signaling, neurotransmitter transport, and dendritic/axonal maturation. For example, expression of DLX5 and DLX6 supports commitment toward GABAergic interneuron identity [39], while upregulation of GRIN1 is indicative of excitatory synaptic priming [40]. These transcriptional findings were further corroborated by microscopy imaging (Figure 2), which revealed neuron-like morphological patterns, including neurite projections, soma thickening, and arborization in treated cells.

Interestingly, gene expression patterns also indicated suppression of markers linked to migratory excitatory neurons, pointing to a transition away from progenitor-like states and toward post-migratory, lineage-stabilized neuronal fates (Figure 4). This shift may reflect decreased cell motility and reduced proliferative capacity along the cortical developmental axis. Although excitatory neuron markers were predominantly enriched, inhibitory markers were also prominently expressed, suggesting a hybrid neuronal identity characterized by the co-expression of both lineage transcription programs (see Supplementary Information).

### 2. Comparative Effects of Neuronal Reprogramming, Astrocytic Differentiation, and H3K27M Knockout Reveal Divergent Therapeutic Outcomes

Our results demonstrate that neuronal reprogramming induces transcriptional phenotype characterized by reduced plasticity and neuronal-like identity markers compared to astrocytic differentiation (DM) or H3K27M knockout (KO). While astrocytic differentiation promoted the expression of glial lineage markers, it also activated genes associated with tumor–immune microenvironment interactions that support tumor progression, including migration, angiogenesis, and extracellular matrix remodeling; traits consistent with a reactive, perivascular-like astrocytic phenotype (Figure 5). Although DM can promote differentiation, its associated migratory programs may preserve or even exacerbate glioma plasticity and invasiveness.

In contrast, H3K27M knockout in serum conditions induced a phenotypic bias toward astrocytic identity while concurrently suppressing neuronal lineage programs. However, we acknowledge that in vivo xenograft models, where K27M-KO delays or abolishes tumorigenesis [38], are essential for validating these in vitro observations. The transcription factor ZNF501, which was strongly associated with the KO condition differential signatures, has been recently identified as a regulator of glioblastoma growth [41] and may represent a potential therapeutic target due to its role in maintaining tumorigenic states. These findings underscore the differential impact of DM and KO conditions on glioma cell fate and highlight the relative advantage of neuronal reprogramming in reducing malignancy-associated plasticity, and potentially a constrained, post-mitotic state for cell fate control.

### 3. Future Directions: Targeted Approaches to Differentiation Therapy in Glioma Reprogramming

While the current drug cocktails promoted neuronal differentiation, they served as valuable discovery tools for identifying key gene signatures that may guide the development of safer, precision-based reprogramming therapies. FID employs GSK3 inhibitors (WNT activators) and BET inhibitors, while FTT includes TGFβ and Rho kinase inhibitors. Our differential expression analysis suggests that the identified gene signatures map onto downstream pathways modulated by these targets, and their mechanisms of action. As previously noted, these combinations activated genes specific to neurodevelopmental programs such as ARX, DLX5, DLX6, BCL11B, WNT7A, NRXN3, NRXN1, and ERBB4—as critical biosignatures of neuronal lineage differentiation (Figure 3B). Interestingly, ERBB3 and WNT7B were downregulated while ERBB4 and WNT7A were upregulated (Figure 3B), though whether this reflects a broader developmental program switch or subtype-specific identity remains to be elucidated, as these genes are commonly expressed across multiple neuronal fate trajectories (See Supplementary Information).

Our findings suggest that ion channels and cell junction-modulating genes such as GRIN1 and NRXN1—upregulated during neuronal reprogramming—may play a key role in restoring disrupted bioelectric networks underlying cellular communication in glioma, thereby contributing to fate stabilization and malignancy suppression (see Supplementary Figures S7–S8, Table S5). Notably, NRXN1 upregulation, observed in our study, has been shown to drive glioblastoma trans-differentiation into neuron-like states [24]. In addition, we propose that ionotropic receptor regulators such as GRIN1 and GRIK1, along with other reprogramming-related DEGs like CXCR4, SEMA3C, and CDH1, may represent precision targets for cellular reprogramming. However, the precise nature of the neuronal subtype identity remains uncertain, as gene expression profiles suggest a potential hybrid identity characterized by both excitatory and inhibitory features. To resolve this ambiguity, single-cell RNA-seq trajectory inference and multiomic integration could help elucidate subcluster heterogeneity, identify partially reprogrammed states, and decode the epigenetic and metabolic mechanisms steering cell fate decisions. Our identified transcriptional signatures should be functionally validated using CRISPR-RNAi screens targeting key regulators of neuronal-like fate commitment. Future work should also incorporate functional assays, such as proliferation, invasion, and xenograft models, to determine whether these transcriptional states correspond to reduced malignant traits.

## Conclusion

This pilot study provides compelling evidence that H3K27M pHGG cell lines (BT245) can be reprogrammed toward neuronal trans-differentiation via pharmacological perturbations, offering a promising alternative to conventional treatment strategies. Overall, this study advances the concept of differentiation therapy as a precision medicine strategy to constrain tumor progression in aggressive glioma ecosystems [18, 42]. We propose that this approach be extended to other high-grade gliomas and epigenetic glioma variants with stalled differentiation, such as H3.1 K27M and G34V/R subtypes [14, 43].While these are early *in vitro* findings, their ability to direct glioma cells toward less malignant, neuronal fates holds translational promise for future patient-centered precision therapies that prioritize patient safety, specificity, and long-term care.

## Declarations

**Funding:** This research was supported by funding from the McGill University William Dawson Scholar Program.

**Competing Interests:** The authors have no relevant competing interests to disclose.

**Ethics Approval:** Only patient-derived cell cultures were used in this study, all of which complied with institutional ethical and biosafety protocol guidelines and certifications. No human participants, identifiable data, or animal experiments were involved.

**Data Availability:** All data associated with this manuscript are available in the NCBI Gene Expression Omnibus (GEO) repository under accession number: GSE298781.

**Author Contributions:** Supervision: J.M. Conceptualization and design: A.U., J.M. Methodology: A.U., C.H., E.B., M.G., J.M. Data analysis: A.U. Writing – original draft preparation: A.U., J.M. Writing – review and editing: A.U., J.M., C.H., E.B.

## Notes

https://www.ncbi.nlm.nih.gov/geo/query/acc.cgi?acc=GSE298781

### Competing Interest Statement

The authors have declared no competing interest.

