## Supplementary Information for "Cellular Reprogramming of H3K27M Pediatric High-Grade Glioma to Neuron-like State"

| GSEA (Biological Processes) | $P_{adj}$ |
|---|---|
| forebrain development | 6.56E-07 |
| axon guidance | 7.64E-07 |
| neuron projection guidance | 7.64E-07 |
| telencephalon development | 8.73E-07 |
| modulation of chemical synaptic transmission | 8.73E-07 |
| Axonogenesis | 8.73E-07 |
| regulation of trans-synaptic signaling | 8.73E-07 |
| axon development | 4.39E-06 |
| central nervous system neuron differentiation | 3.84E-05 |
| neuron projection extension | 4.13E-05 |

**Table S1.** GSEA Enrichment of upregulated DEGs top 10 pathways for All Treatments DEGs (log FC >2 and p<0.05) ranked by log2 fold-change using the FGSEA R package and Gene Ontology (GO) Biological Processes Database.

As a complementary analysis to Figure 3B in the Main Text, which showed gene set enrichment of neuronal differentiation pathways induced by reprogramming cocktails, we further examined functional enrichment using a web-based tool, g: Profiler, to validate and expand upon these findings. g: Profiler cross-validated the fgsea-identified genes and examined the functional impact of significantly upregulated DEGs to confirm their neuronal differentiation potential (Table S2). While Figure 3B results consist of GSEA via genes ranked by log-fold change, g: Profiler performs overrepresentation analysis by identifying significantly enriched biological terms and pathways from the provided gene list of all upregulated DEGs. The top enriched GO term, "synapse" (padj = $3.901 \times 10^{-13}$), underscores a profound induction of synaptogenesis and enhanced synaptic activity. Related critical terms, such as "neuron projection" (padj = $3.422 \times 10^{-9}$) and "axon development" (padj = $6.589 \times 10^{-9}$), emphasize structural neuronal changes, including dendritic branching, axonal elongation, and improved neuronal connectivity, essential for mature neuronal functionality.

Moreover, enriched GO categories such as "post-synapse" (padj = $3.368 \times 10^{-5}$) and "GABA-ergic synapse" (padj = $1.164 \times 10^{-3}$) suggest specialization toward functional neuronal subtypes, particularly inhibitory neuronal networks, crucial for stable neuronal circuit integration. Genes driving these GO categories include key neuronal markers such as GRIN1, GRIK1, CALB2, SLC32A1, SLC1A2, and NRXN1, known to regulate synaptic plasticity and neurotransmission dynamics. These pathways demonstrate that the pharmacological interventions strongly induce neuronal differentiation programs in BT245 glioma cells, redirecting their phenotype from highly malignant, proliferative states toward hybrid neuronal identities (i.e., with excitatory and inhibitory neuronal traits). Such transcriptional reprogramming highlights the therapeutic promise of differentiation therapy in mitigating pediatric glioma aggressivity, by constraining or re-directing phenotypic (malignant) plasticity.

| GO Term Name | Padj |
|---|---|
| synapse | $3.901 \times 10^{-13}$ |
| neuron projection | $3.422 \times 10^{-9}$ |
| cell junction | $4.144 \times 10^{-9}$ |
| axon | $6.589 \times 10^{-9}$ |
| somatodendritic compartment | $4.665 \times 10^{-7}$ |
| presynapse | $6.233 \times 10^{-7}$ |
| cell projection | $2.820 \times 10^{-6}$ |
| plasma membrane bounded cell projection | $6.273 \times 10^{-6}$ |
| neuron to neuron synapse | $8.246 \times 10^{-6}$ |
| postsynapse | $3.368 \times 10^{-5}$ |
| dendrite | $8.961 \times 10^{-5}$ |
| dendritic tree | $9.409 \times 10^{-5}$ |
| cell body | $1.097 \times 10^{-4}$ |
| cell periphery | $1.140 \times 10^{-4}$ |
| neuron projection terminus | $1.706 \times 10^{-4}$ |
| glutamatergic synapse | $3.019 \times 10^{-4}$ |
| neuronal cell body | $4.548 \times 10^{-4}$ |
| synaptic membrane | $5.941 \times 10^{-4}$ |
| asymmetric synapse | $6.408 \times 10^{-4}$ |
| axon terminus | $6.912 \times 10^{-4}$ |
| GABA-ergic synapse | $1.164 \times 10^{-3}$ |

**Table S2.** g: Profiler Gene Set Enrichment Analysis of Upregulated DEGs ($p_{adj} < 0.05$, $logFC > 2$) in treatment samples show activation of neuronal markers and neuronal differentiation programs.

| Term Name | Adjusted P-Value | Associated Genes |
|---|---|---|
| Forebrain development | 3.01E-05 | BCL11B, DLX6-AS1, ARX, KIF26A, ATP1A2, KIRREL3, SLC32A1, CDH1 |
| Brain development | 0.000176566 | BCL11B, DLX6-AS1, ARX, KIF26A, ATP1A2, GRIN1, KIRREL3, SLC32A1, CDH1 |
| Head development | 0.000303748 | BCL11B, DLX6-AS1, ARX, KIF26A, ATP1A2, GRIN1, KIRREL3, SLC32A1, CDH1 |
| Telencephalon development | 0.001310549 | BCL11B, ARX, KIF26A, ATP1A2, KIRREL3, SLC32A1 |
| Central nervous system development | 0.002842217 | BCL11B, DLX6-AS1, ARX, KIF26A, ATP1A2, GRIN1, KIRREL3, SLC32A1, CDH1 |
| Anterograde trans-synaptic signaling | 0.033045806 | ATP1A2, CALB2, GRIN1, SCGN, CLSTN2, SLC32A1, CDH1 |
| Chemical synaptic transmission | 0.033045806 | ATP1A2, CALB2, GRIN1, SCGN, CLSTN2, SLC32A1, CDH1 |
| Synaptic signaling | 0.042942718 | ATP1A2, CALB2, GRIN1, SCGN, CLSTN2, SLC32A1, CDH1 |
| Synapse | 0.00089526 | IQSEC3, ATP1A2, CALB2, GRIN1, SYNPR, SCGN, KIRREL3, CLSTN2, SLC32A1, CDH1 |
| Dendrite | 0.001501629 | ATP1A2, CALB2, GRIN1, SCGN, KIRREL3, CLSTN2, SLC32A1 |
| Dendritic tree | 0.001533505 | ATP1A2, CALB2, GRIN1, SCGN, KIRREL3, CLSTN2, SLC32A1 |
| Axon terminus | 0.002066515 | CALB2, GRIN1, SCGN, SLC32A1 |
| Neuron projection | 0.003001883 | BCL11B, ATP1A2, CALB2, GRIN1, SYNPR, SCGN, KIRREL3, CLSTN2, SLC32A1 |
| Neuron projection terminus | 0.003395887 | CALB2, GRIN1, SCGN, SLC32A1 |

Table S3. G: Profiler Analysis of the statistically most significant 30 DEGs for Combined Neuronal Differentiation Treatments (FTT and FID Cocktails). These findings suggest the BT245 cell fates are transitioning towards neocortical (telencephalon and forebrain) neuron-like states.

**Astrocytic Differentiation Results in Partial Lineage Shift and Vascular Niche-Associated Traits**

As shown in Figure S1A, PCA mapping of bulk RNA-seq transcriptomes revealed distinct transcriptional profiles between BT245 cells cultured in stem-cell proliferation media (SCM) as controls, and those treated with astrocytic serum-differentiation media (DM). For this analysis across 56,226 genes, we identified 570 downregulated and 2,641 upregulated genes using the same stringent criteria as in previous analyses. A clear separation is observed between control and astrocytic differentiation samples along the first principal component axis. These results suggest a robust transcriptional response to astrocytic differentiation conditions, confirming that serum-based DM induce distinct astrocytic lineage programs compared to standard proliferation conditions. Furthermore, as shown in Figure S1B, red-labeled genes (e.g., *LUM, CFH, COL1A1*) represent the top upregulated DEGs, many of which are associated with extracellular matrix remodeling and angiogenesis, characteristic of invasive glioma phenotypes. Blue-labeled genes (e.g., *GABRA5, IGF2, MKX*) are among the top downregulated DEGs, reflecting the suppression of neuronal signaling and differentiation pathways in the astrocytic DM condition. Among the top 50 significantly upregulated DEGs in the astrocytic DM condition were GREM1 and BMP5, both involved in developmental signaling, along with astrocytic markers such as AQP4. These genes reflect activation of a differentiation program biased toward perivascular and glial lineages. The overall DEG profile suggests enhanced extracellular matrix remodeling, vascular interaction, and migratory potential.

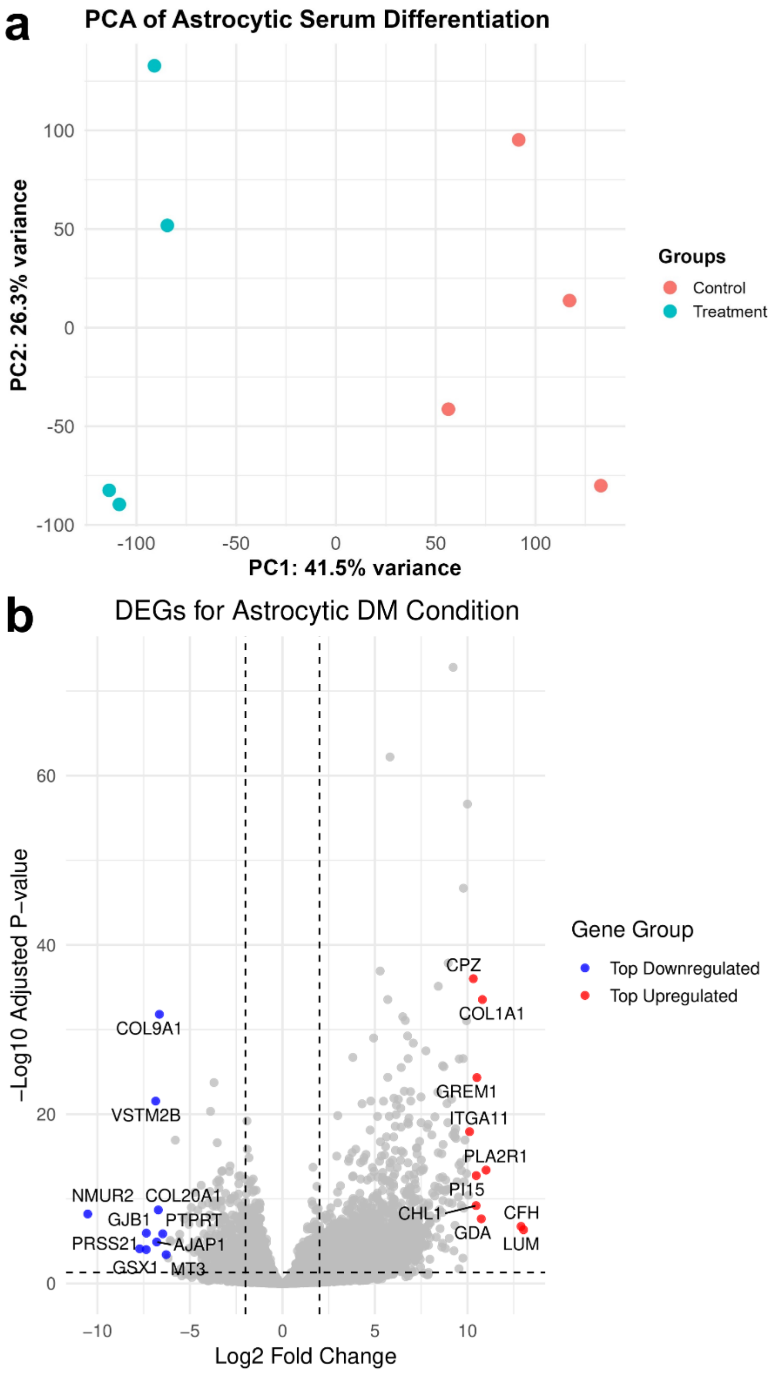


**Figure S1. A) PCA Clustering of Control BT245 vs. Astrocytic Serum Differentiation Media Treatment.** The K27M BT245 cell lines were cultured in neural stem-cell proliferation media (SCM) as controls and in astrocytic serum-differentiation media (DM) to promote astrocytic differentiation, two weeks before bulk RNA sequencing. B) Top 10 upregulated and downregulated genes in the Astrocytic DM condition (Figure S1B): Volcano plot highlighting the most significantly upregulated and downregulated genes ($padj < 0.05$) based on log2 fold-change. Genes involved in ECM remodeling, angiogenesis, and tumor-promoting pathways are prominently induced, while neural differentiation markers are suppressed, indicating a transcriptional program associated with invasion and perivascular niche adaptation.

Gene expression profiling under astrocytic DM conditions revealed transcriptional transitions in BT245 glioma cells, encompassing both neuronal differentiation pathways and vascular niche-cell interactions (Figure S2). Despite the astrocytic differentiation cues provided by the media, upregulated DEGs (Figure S2A) still prominently included markers associated with the neuronal TLE4 subtype, suggesting residual neurogenic potential within the treated glioma cells or overlapping differentiation programs in their preferred cell identity, cell fate bias, or lineage commitment [44]. Further, focused analysis of the top 100 upregulated DEGs (Figure S2B) revealed significant alignment with pericyte (Per-1) and vascular leptomeningeal cell (VLMC-1) signatures, which are key cellular components of perivascular niche construction. VLMC-1 cells, characterized by fibroblast-like traits, function at the critical interface between astrocytes and the blood-brain barrier (BBB) and harbor inherent neurogenic properties [45]. The prominent expression of these niche-associated markers indicates a potential hijacking of perivascular differentiation routes by the astrocytic DM-exposed glioma cells, potentially facilitating their survival and persistence near BBB interfaces and allowing invasive niche construction capabilities. As such, these transcriptional changes could potentially enhance glioma malignancy and invasiveness, rather than steering cell fates towards stable (terminal) astrocytic differentiation, though these predictions require validation through functional assays

Conversely, downregulated DEGs under astrocytic DM conditions predominantly indicated suppression of mesenchymal stem cell (MSC-1/2) traits (Figure S2C). This pattern is characterized by reduced expression of genes linked to migratory and invasive phenotypes, suggesting a transcriptional shift away from aggressive mesenchymal states. The continued partial expression of neuronal and perivascular niche-cell signatures implies that glioma cell plasticity is not entirely suppressed. Rather, these cells may retain adaptive behaviors, such as phenotypic plasticity, navigating differentiation pathways in response to varying microenvironmental pressures, highlighting the complexity and partial commitment of glioma cells under astrocytic differentiation conditions.

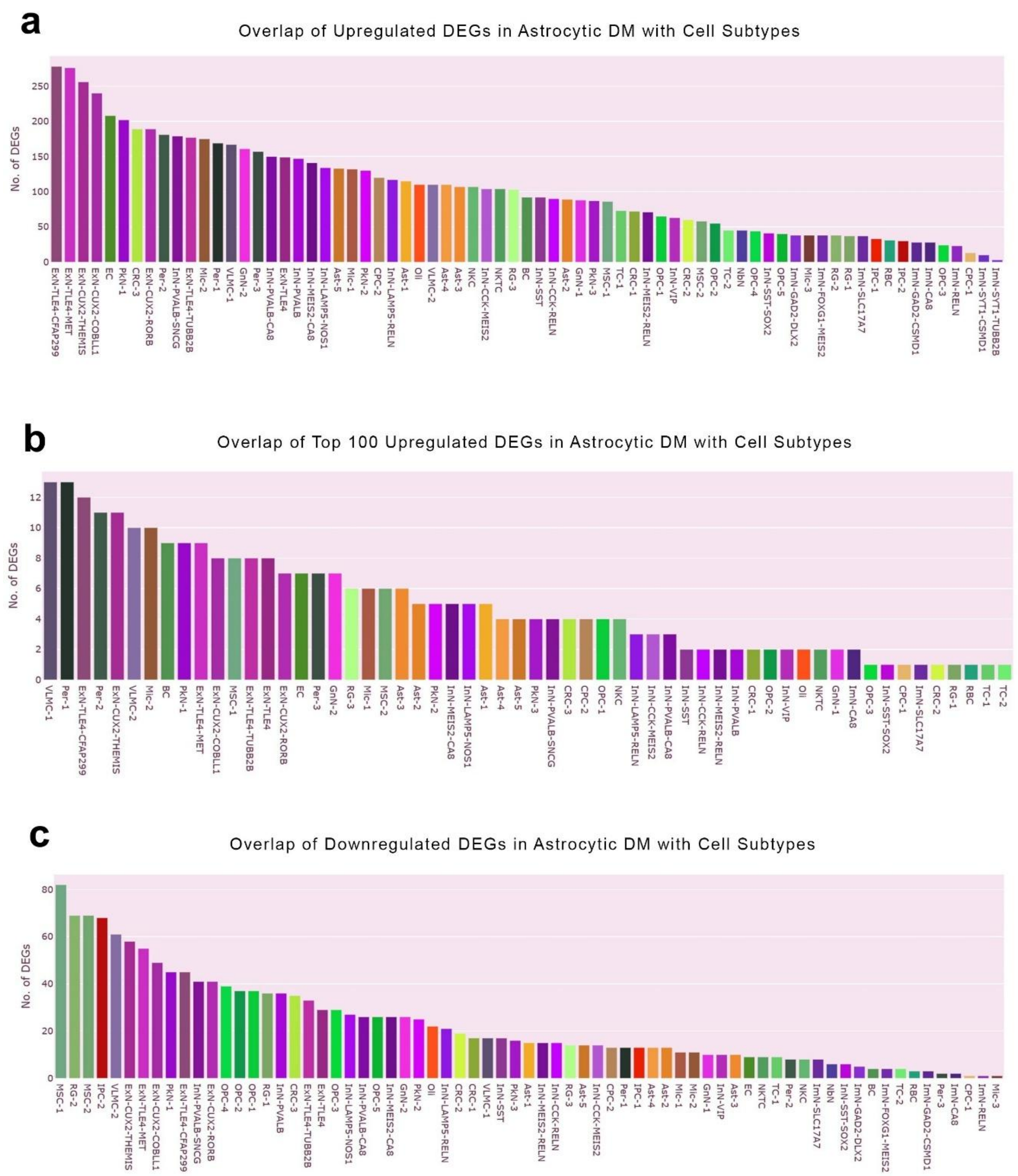


**Figure S2. Cell Type Analysis of DEGs in DM condition.** A) Upregulated DEGs ($p<0.05$, $logFC > 2$) suggest that TLE4 subtype excitatory neuron markers are still the most highly expressed in DM astrocytic differentiation conditions. B) STAB-2 analysis of only the top 100 upregulated DEGs shows that the Per-1 (pericytes) and VLMC-1 (vascular leptomeningeal cells) cell types are the best predictors of phenotypic expression. C) Downregulated DEGs in astrocytic DM conditions suggest signatures of MSC-1/2 (mesenchymal stem cells) are most suppressed.

**Astrocytic Differentiation Activates Perivascular and Migratory Programs Associated with Tumor Invasion**

The g: Profiler functional enrichment analysis shown in Table S4 of upregulated DEGs from the astrocytic serum differentiation media (DM) treatment highlighted prominent activation of pathways linked to extracellular matrix remodeling, and cancer invasion. For instance, the robust enrichment of terms such as "cell adhesion" (padj = $5.009 \times 10^{-39}$), "cell migration" (padj = $1.225 \times 10^{-30}$), "angiogenesis" (padj = $6.139 \times 10^{-23}$), and "blood vessel development" (padj = $2.798 \times 10^{-29}$) highlights the strong transcriptional induction of genes facilitating cellular interactions with the perivascular niche and BBB. Specifically, genes like AQP4, a water channel marker predominantly expressed by astrocytes at the BBB interface and critical for astrocytic migration, illustrate how astrocytic DM conditions may activate transcriptional programs that promote glioma cell survival, invasion, and phenotypic plasticity within vascularized immune-tumor microenvironments. These findings further imply that astrocytic DM not only drives glioma cells toward astrocytic lineage features but also potentially facilitates tumor cell persistence, niche construction, and invasiveness.

| GO Term Name | Padj |
| --- | --- |
| extracellular matrix binding | $4.622 \times 10^{-8}$ |
| cell adhesion | $5.009 \times 10^{-39}$ |
| circulatory system development | $3.121 \times 10^{-35}$ |
| cell migration | $1.225 \times 10^{-30}$ |
| extracellular matrix organization | $1.658 \times 10^{-29}$ |
| extracellular structure organization | $2.143 \times 10^{-29}$ |
| blood vessel development | $2.798 \times 10^{-29}$ |
| blood vessel morphogenesis | $3.915 \times 10^{-25}$ |
| regulation of cell adhesion | $1.042 \times 10^{-24}$ |
| locomotion | $3.606 \times 10^{-23}$ |
| angiogenesis | $6.139 \times 10^{-23}$ |

| | |
|---|---|
| positive regulation of cell communication | $3.139 \times 10^{-22}$ |
| cell-cell adhesion | $3.904 \times 10^{-21}$ |

**Table S4. g: Profiler Analysis of Upregulated DEGs from Astrocytic DM treatment shows Migratory phenotypes. G: Profiler analysis of** upregulated DEGs confirms the results from the STAB-2 cell type assignment algorithm, i.e., pericytes and VLMC-1-like parenchymal niche cell phenotypes.

### H3K27M Knockout (KO) Preserves Plastic Cellular Identity, While Serum Differentiation Enhances Cell Fate Bias Towards Astrocytic-Like Reprogramming

For the K27M knockout (KO) condition, DESeq2 analysis across 56,226 genes identified 84 upregulated and 59 downregulated genes using the same stringent thresholds (absolute log2 fold change ≥ 2, Benjamini-Hochberg corrected p-value < 0.05). PCA clustering revealed a clear divergence in transcriptional profiles between K27M knockout (KO) conditions under different culture environments (Figure 6). H3K27M KO cells maintained in stem-cell proliferation media (SCM) clustered closely with untreated BT245 controls, indicating that KO alone does not alter the underlying plastic glioma state. This suggests that in the absence of additional differentiation cues or signals, the KO cells preserve an undifferentiated transcriptomic profile. However, when K27M KO cells were exposed to astrocytic differentiation media (DM) (SerDiff_KO), they formed a distinct cluster, separated along PC1 in PCA space, which accounts for 46% of total variance, and PC2, showing an additional 26%.

Furthermore, DESeq2-based differential expression analysis revealed that out of the 59 significantly downregulated genes in the K27M KO condition, GABRA5—which encodes a subunit of the $GABA_A$ receptor critical for inhibitory synaptic signaling—was suppressed, while GRM5, a gene encoding a metabotropic glutamate receptor, was among the 84 upregulated DEGs (Figure 6B). This reciprocal transition suggests a loss of GABAergic neuron-like properties and a potential redirection toward glutamatergic signaling following K27M depletion. Several additional gene signatures shown on the volcano plot in Figure 6B highlight a shift in differentiation programs following H3K27M knockout. Among the significantly upregulated genes are CD74, CIITA, and TNFRSF11A, which are involved in immune-inflammatory signaling, such as NF-κB pathway activation, in KO conditions. SLC16A3 and SLC47A2, which encode lactate transporters and multidrug extrusion proteins, respectively, reflect metabolic remodeling along with the upregulation of IGFBP7, a known modulator of IGF signaling and senescence. Conversely, among the downregulated DEGs, in addition to GABRA5, suppression of SLC2A12, a glucose transporter, may indicate altered metabolic programs; and ARHGAP26 encoding a Rho GTPase.

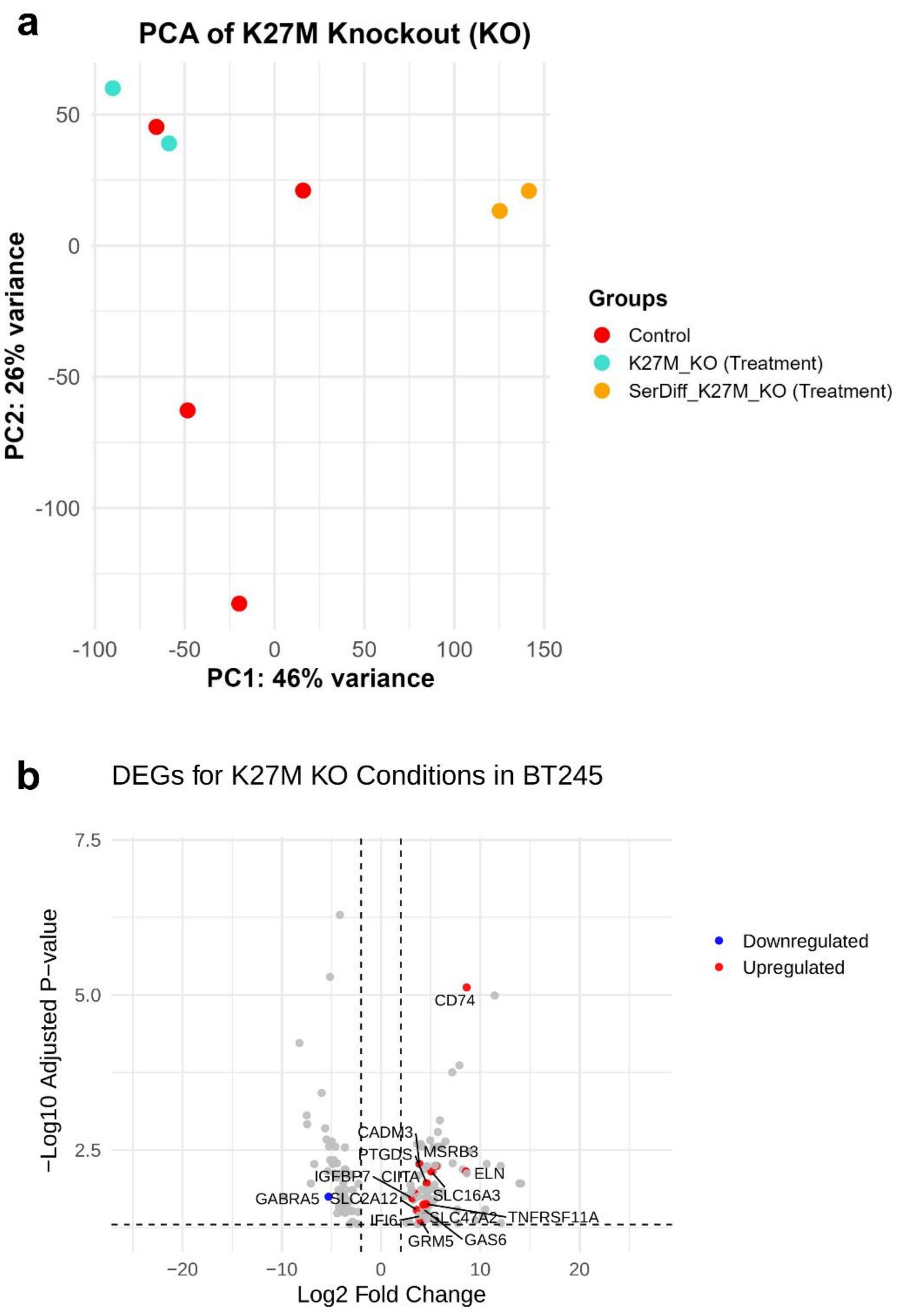


**Figure S3. A) PCA Clustering of H3K27M Knockout (KO) Conditions.** KO samples (in Turquoise) maintained in stem cell media (SCM) cluster tightly with control BT245 glioma cells (in Red), indicating that KO alone does not induce major transcriptomic reprogramming. This suggests that in the absence of external differentiation cues, the K27M KO cells retain a glioma-like, stem cell-associated identity. In contrast, K27M KO cells cultured in astrocytic serum differentiation media (DM) (shown in yellow) form a distinct cluster, highlighting a synergistic reprogramming effect driven by the combination of epigenetic de-repression and astrocytic serum-induced lineage-specific signals. **B) Volcano Plot of Differentially Expressed Genes**

**(DEGs) in K27M KO Conditions.** The plot highlights selected **protein-coding genes** with significant upregulation (red) or downregulation (blue) based on stringent thresholds ($|\log_2$ fold change$| > 2$ and Benjamini-Hochberg corrected p-value $< 0.05$, DESeq2 analysis).

**H3K27M Knockout in DM Conditions Promotes Astrocytic Differentiation and Suppresses Neuronal Identity**

The separation observed in the PCA analysis of Figure S3A is further substantiated by Figure S4A, which shows that upregulated DEGs in the KO+DM condition are strongly enriched for markers of perivascular astrocytes, microglia, and endothelial cells. Figure S3B shows that upregulated genes included CD74, CIITA, TNFRSF11A, and SLC16A3, which are associated with immune signaling, metabolic remodeling, and senescence pathways. Notably, GABRA5, a key marker of GABAergic neuronal identity, was significantly downregulated, indicating suppression of some neuronal programs post-KO. Key astrocytic and niche-associated markers, including COLEC12, ELN, JUP, CPAMD8, TMEM59L, and PTGDS, are elevated, many of which are transcriptionally coordinated under the influence of ZNF501, a transcription factor associated with glial specification, as revealed by g: Profiler analysis. These transcritpional changes suggest a cell fate bias towards an astrocytic lineage.

Meanwhile, Figure S4B reveals that neuronal lineage markers are markedly suppressed in KO+DM samples, confirming that neuronal differentiation programs are actively downregulated under these differentiation conditions. This indicates that the H3K27M mutation may serve as a molecular barrier to astrocytic commitment, aligning with current theories of the developmental blockade or stalled differentiation of gliomas along their neurodevelopmental hirarchies [14-16], and its removal facilitates glial differentiation under the controlled environmental stimuli.

Thus, K27M-KO appears to promote an astrocytic phenotype while suppressing neuronal programs, particularly when combined with serum differentiation signals. While this astrocytic transition may suppress malignant features such as plasticity seen in DM alone conditions (Figure S2), it also limits the potential for transdifferentiation toward less invasive, post-mitotic neuronal fates (although this requires validation through functional assays). These findings suggest a critical dichotomy between astrocytic and neuronal reprogramming strategies in glioma cell fate control, with the latter potentially offering a more stable and therapeutically favorable outcome in the context of differentiation therapy.

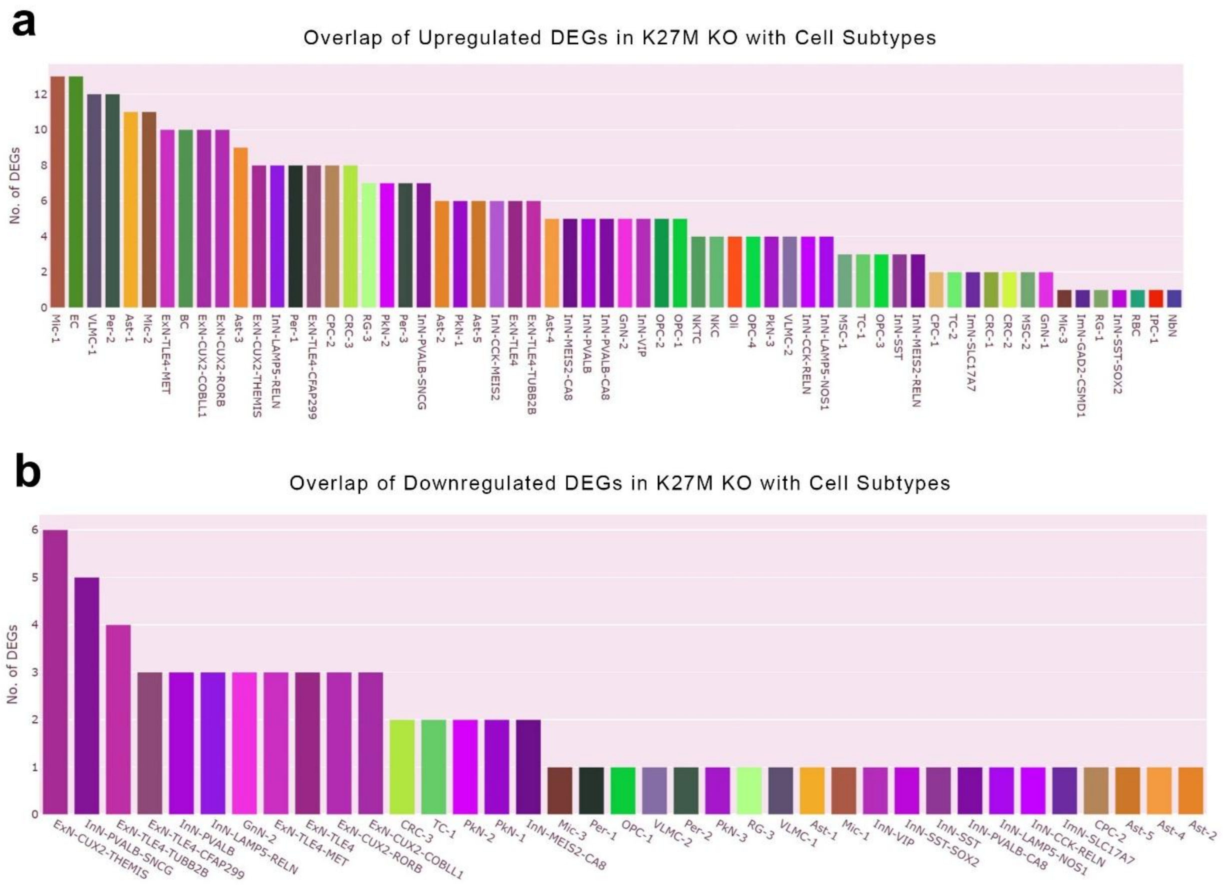


**Figure S4. Cell Type Assignment with DEGs in K27M KO conditions.** A) Upregulated DEGs (log2FC > 2, $p < 0.05$). B) Downregulated DEGs (log2FC < -2, $p < 0.05$). The upregulated genes resemble the results shown in Figure 5B, for astrocytic DM. The upregulated DEGs most overlap with microglia subtypes, endothelial cells, and perivascular astrocytes, while most of the downregulated genes overlap with neuronal phenotypes.

**Network Analysis of Neuronal Trans-differentiation DEGs Reveals Ion channels and Bioelectric Networks as Targets for Cell Fate Reprogramming**

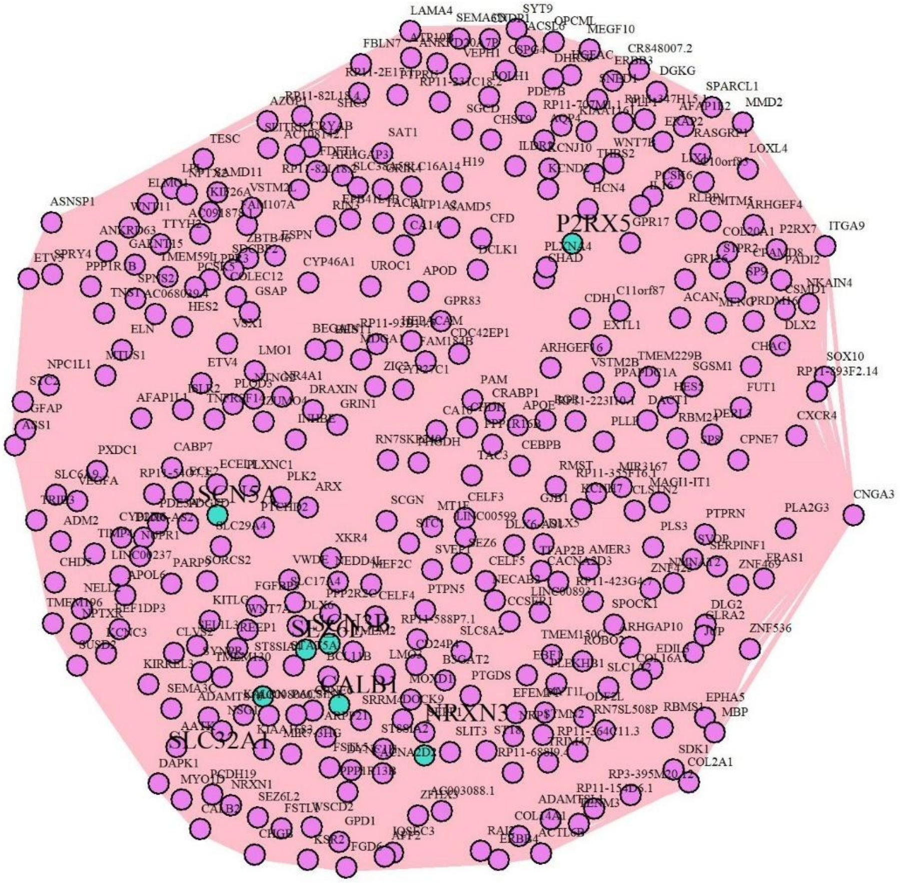


**Figure S7. Spearman Correlation Network between DEGs in Samples with Neuronal Trans-differentiation Treatments (FID and FTT combined).** The Spearman correlation was used as a simple network inference metric to identify centralities in the neuronal reprogramming treatment groups. Some network centralities (i.e., key regulators of gene regulatory network dynamics) are colored in turquoise. The nodes with the highest eigenvector centrality (i.e., nodes with the highest connectivity and outgoing information flow) included BC11B, SEMA3C, NRXN3, NRXN1, WNT7A, SLC1A2, SLC32A1, SCN3B, SCN5A, CALB1, DLX6, FGFBP3, MOXD1, and P2RX5**.** All identified network centrality markers promote neuronal differentiation, mostly by altering the activity patterns of voltage-gated ion channels/transporters, synaptic/neurodevelopmental genes, and differentiation signaling genes.

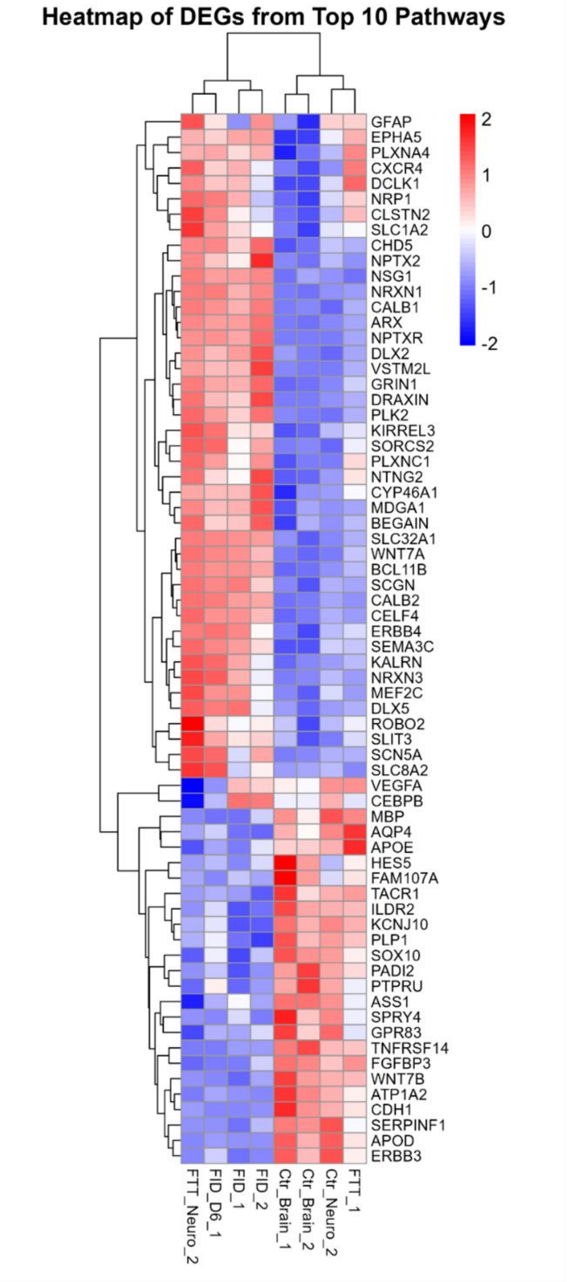


**Figure S8.** Heatmap showing the top DEGs from the top 10 GO: BP enriched pathways identified by fgsea analysis in Figure 3B. Expression patterns reflect neuronal differentiation signatures across treatment and control conditions.

#### Network Centrality Measures Highlight Key Cell Fate Regulators and Potential Therapeutic Targets for Neuronal Differentiation

Network centrality measures provide quantitative insights into the complex dynamics and topology (structure) of gene interaction networks in neuronal differentiation. Closeness centrality identifies nodes that efficiently spread information across the network, while betweenness centrality highlights key bridge nodes that regulate information flow between modular compartments. Degree centrality points to nodes with many direct connections, while eigenvector centrality indicates highly influential nodes connected to other central 'hubs' (i.e., authoritative clusters). Together, these metrics offer quantitative tools for identifying critical regulators of cell fate transitions within the network. P2RX5 emerged as the top-ranking gene for closeness and betweenness centralities, followed by NPC1L1. P2RX5, a ligand-gated ion channel (purinoceptor for ATP), is central to neuronal signaling. CDH1, NRXN1/3, BC11B, PPP2R2C, SYNPR, ARPP21, and SEMA3C, among other network signatures, were observed as the highest eigenvector, degree, and BDM perturbation signatures. These findings suggest neuronal-like identity markers, while DLX6 (or DLX6-AS1), WNT7A, HES5, and FGFBP3 indicate neurodevelopmental (morphogenetic) programs coordinating the patterning processes. Gene set enrichment analysis (g:Profiler) further supports these findings, indicating these genes map to pathways associated with neuronal lineage and development.

Additionally, the g: Profiler results emphasized enriched GO terms with significant p-values within the Spearman network. The most prominent term, neuron projection morphogenesis (GO:0048812; $p = 8.864 \times 10^{-6}$), highlights its critical role in the DEGs. Other key terms, such as synapse organization ($p = 4.176 \times 10^{-5}$), modulation of chemical synaptic transmission ($p = 5.172 \times 10^{-3}$), and voltage-gated sodium channel activity involved in action potential ($p = 3.409 \times 10^{-2}$), underscore essential processes in neuronal development and signaling. These findings highlight central plasticity regulators and pathways driving neuronal-like lineage differentiation and thus, potential therapeutic targets for cell fate reprogramming.

#### BDM Reveals Key Regulators of Neurogenesis and Synapse Formation in Network Perturbation Analysis

The Block Decomposition Method (BDM) measures algorithmic complexity, the minimal description length of a complex system, by quantifying the structural organization and regularities of a network [36]. It provides computational approach to identifying causal information dynamics by assessing how perturbations (i.e., gene node or link deletions on the Spearman network) disrupt the network topology and stability. Thus, BDM-based perturbation reveals which network signatures are critical for maintaining the network's functional robustness and causal structure, offering actionable insights into the mechanisms driving the complex system's patterns of behavior (dynamics). BDM perturbation analysis identified critical regulators of neurogenesis and synapse formation. Perturbation results highlighted significant GO terms such as synapse ($p = 8.459 \times 10^{-11}$), cell junction ($p = 1.370 \times 10^{-8}$), cell projection ($p = 5.184 \times 10^{-6}$), and neuron projection ($p = 8.700 \times 10^{-6}$), along with Reactome pathways including Neurexins and Neuroligins ($p = 1.075 \times 10^{-2}$) and Neuronal System ($p = 1.493 \times 10^{-2}$). These results further support the activation of biological processes underlying neuronal differentiation by the FTT/FID cocktail treatments. Key synaptic signatures identified by the highest BDM perturbation shifts included IQSEC3, KIRREL3, CDH1, NRXN3, PTPRN, NRXN1, KCNC3, KCNJ10, SLC1A2, SLC32A1, SLC29A4, P2RX5, SEZ6L, SYNPR, and WNT7A, all of which are critical for neurotransmission and synaptic organization.

| **BDM Change** | **Eigenvector** | **Degree** |
|---|---|---|
| DLX6-AS1 | BCL11B | CDH1 |
| IQSEC3 | SEMA3C | PLLP |
| KIF26A | ARPP21 | APOD |
| ECEL1 | LMO3 | CA10 |
| ARPP21 | PPP2R2C | PLA2G3 |
| ZNF469 | SYNPR | CRABP1 |
| TNFRSF14 | FGFBP3 | APOL6 |
| SCGN | KIRREL3 | PAM |
| KIRREL3 | KITLG | RGR |
| KITLG | SLC32A1 | FUT1 |
| CDH1 | SRRM4 | RLBP1 |
| RP11-93B14.5 | DLX6 | P2RX5 |
| CRYAB | B3GAT2 | PARP9 |
| ESPN | CPNE6 | CHDH |
| CPNE6 | KIAA1683 | APOE |
| EFEMP1 | CELF4 | VSX1 |
| ZIC2 | TMEM2 | UROC1 |
| LPPR3 | SCN3B | MBP |
| KIAA1683 | WNT7A | MTUS1 |
| TMEM2 | MOXD1 | GPR17 |
| SAMD11 | NRXN3 | GJB1 |
| PCDH19 | PTPN5 | AATK |
| MOXD1 | MIR7-3HG | ZFHX3 |
| NRXN3 | DYNC1I1 | CMTM5 |
| PTPRN | ADAMTS10 | H19 |
| ISLR2 | KALRN | SP8 |
| MIR7-3HG | RP11-588P7.1 | HES5 |
| SVEP1 | SLC8A2 | TMEM229B |
| NUPR1 | REEP1 | VSTM2B |
| FGD6 | SEL1L3 | CPNE7 |

**Table S5.** Top 30 Centrality Measures using BDM Perturbation Analysis on Spearman Network from Neuronal Differentiation DEGs. NRXN1 was also seen among the top 35 network centrality signatures.

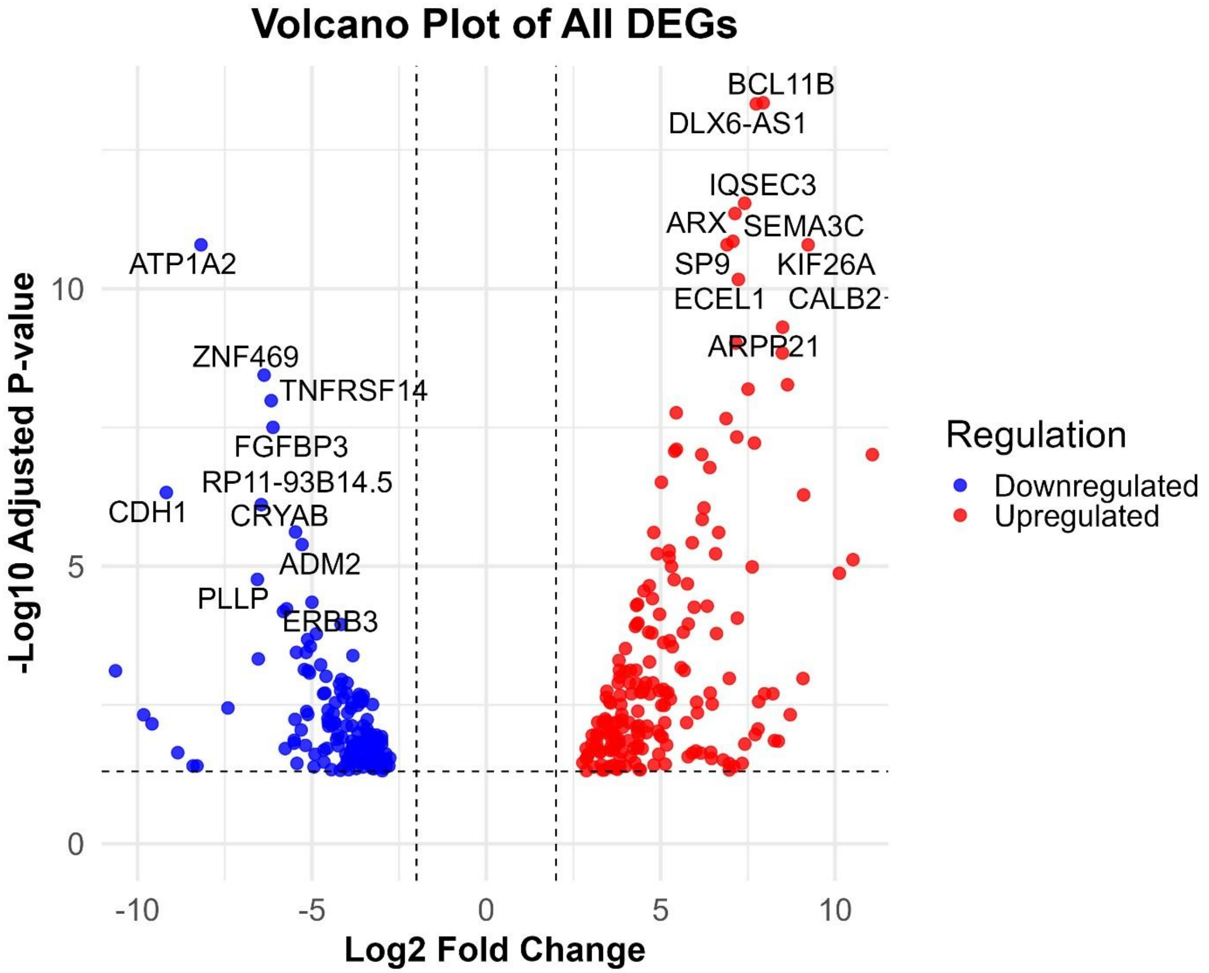


**Figure S9.** Volcano plot displaying DEGs across all treated samples (FID and FTT; Early and Late) combined versus controls, from the comparison shown in Figure 3B of the main text, highlighting top 30 significantly upregulated (red) and downregulated (blue) genes (|log2FoldChange| > 2, padj < 0.05, Benjamini-Hochberg corrected).

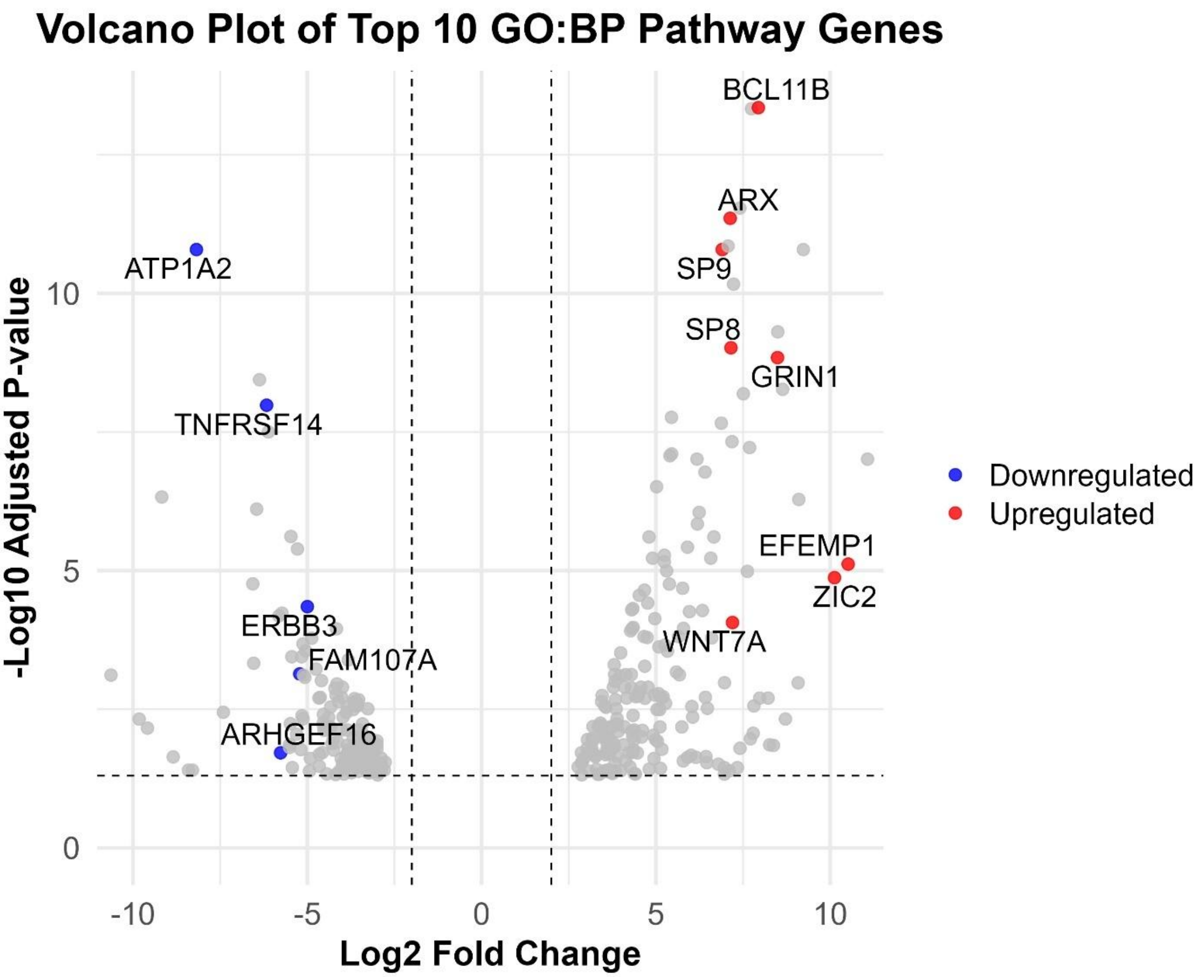


**Figure S10. fgsea on Full Gene Expression Profile Reveals Neuronal Differentiation Pathway Enrichment.** To systematically characterize the gene expression programs that are affected by the neuronal differentiation regimen, we performed fast gene set enrichment analysis (fgsea). Figure 3B (Main Paper) shows the results of top 10 significantly enriched GO: Biological Process (GO: BP) pathways fgsea performed on only the DEGs identified using DESeq2, ranked by log2 fold-change. Herein, Figure S10 displays the top 10 significantly enriched GO: BP pathways identified using fgsea on the entire gene expression profile, ranked by log2 fold-change. This analysis revealed strong enrichment for neurodevelopmental and morphogenetic processes, including *appendage morphogenesis*, *embryonic development*, *forebrain development*, and *telencephalon development*. Key genes contributing to these pathways include ZIC2, GRIN1, ERBB4, BCL11B, and ARX. Additional influential genes linked to these pathways (not shown) include SEMA3C, DLX5, DLX6, SLC32A1, TBX3, FGF9, FGF13, NR2E1, LHX2, DRAXIN, CXCR4, SCN5A, FOXC2, and GREM1.

### Additional Discussions and Suggested Interpretations

#### 1. Functional Roles of Top DEGs in Neuronal Subtype Specification and Bioelectric Modulation

Many of the DEGs highlighted a cell fate bias towards neuronal-like identities. Upregulated DEGs, including DLX5 and DLX6, are functionally implicated in promoting GABAergic differentiation, critical for inhibitory circuit specification [39]. CALB2, a calcium-binding protein, plays key roles in inhibitory neuronal circuits and synaptic plasticity [46]. GRIN1, encoding a subunit of the NMDA receptor, is essential for glutamatergic neurotransmission and synaptic plasticity [40]. SLC32A1 (vesicular GABA transporter) and SLC1A2 (glutamate transporter) reflect transcriptional priming of both inhibitory and excitatory neurotransmitter systems, supporting the emergence of a hybrid excitatory/inhibitory neuronal phenotype. Similarly, FGFBP3 elevation suggests activation of FGF signaling pathways involved in neurodevelopmental progression. These findings support a model in which the reprogramming cocktails initiate partial lineage re-specification into multiple neuron-like identities while repressing glioma-related bioelectric instability, through the regulation of genes like KCNJ10, GRIN1, NRXN1, and NRXN3, all involved in ion channel function and synaptic modulation [24, 47-51]. We observe co-expression of excitatory and inhibitory neuronal markers, indicative of a hybrid neuronal identity.

#### 2. Functional Interpretation of Key Genes Enriched in fgsea Pathways (Related to Figure 3B)

To further interpret the pathways enriched in Figure 3B of the main manuscript, which displays the top Gene Ontology: Biological Process (GO: BP) terms from fgsea analysis performed on the entire log2FC-ranked gene expression profile, we examined the individual genes contributing most to these neurodevelopmental pathways. These genes not only exhibited the highest levels of upregulation in response to the treatments (Figure 3B) but also played central regulatory roles in synaptogenesis, axonal projection, and forebrain specification (Figure 3B). For instance, ARX encodes a transcription factor critical for cortical neuron specification, and neuronal patterning processes [52]. SEMA3C has been demonstrated as a key driver of Wnt pathway activation and resistance in glioblastoma [53]. Notably, WNT7A, known to promote neuronal differentiation, promotes the transition of radial glia into intermediate neural progenitors along a neuronal lineage, sustaining neurogenesis in the developing cortex [54]. Meanwhile, WNT7B, associated with progenitor maintenance and glial identity, is downregulated (Figure 3B). WNT7B overexpression disrupts expression of key pro-neural transcription factors, thereby impairing the maturation of intermediate neuronal pools from radial glia [55, 56]. These findings align with emerging evidence from single-cell lineage tracing studies revealing intermediate progenitor cell trajectories from radial glia (neural stem cells) in H3K3K27M gliomas [15]. GRIN1 and SLC32A1 are key regulators of excitatory and inhibitory neurotransmission, respectively, consistent with the observed shift toward differentiated neuronal-like identities [40, 57]. The upregulation of both inhibitory and excitatory neuronal markers suggests a mixed, partially reprogrammed intermediates or hybrid neuronal identity, with potential for differentiation into multiple neuronal subtypes (Figure 3B).

Downregulated log2FC-ranked, fgsea-linked genes include TACR1, linked to inflammatory signaling and cortical GABAergic neurons [58]; P2RX7, involved in pro-inflammatory responses across the nervous system [59]; PARP9, implicated in inflammation and proliferation pathways; and HES5, known for the transitions of neocortical neural stem cells and progenitors during development [60]. These findings confirm that the drug-induced transcriptional reprogramming engages a coordinated set of neurodevelopmental programs driving the observed cell fate transition.

#### 3. Experimental Rationale for DM and KO Reprogramming Conditions

As shown in Figure 5 of the main paper, the DM condition was designed to promote astroglial lineage differentiation [37, 38], whereas the KO condition aimed to relieve epigenetic repression and restore stalled developmental programs [8, 37]. Detailed analyses of these reprogramming strategies and their gene expression changes are provided in the Supplementary Information: Figures S1 and S2 for DM, and Figures S3 and S4 for the KO condition. Notably, five genes (COLEC12, CPAMD8, ELN, JUP, PTGDS) were consistently upregulated across all three treatments. While this overlap was observed directly from DEG lists, statistical assessment (e.g., Jaccard similarity) may further validate whether such convergence exceeds random expectation. Due to the stringent log2FC cutoff, TMEM59L emerged as the only overlapping DEG between the ND and KO conditions.

#### 4. Suppression of Glial and Malignancy-Associated Signatures During Neuronal Reprogramming

Neuronal trans-differentiation was associated with marked downregulation of markers related to glial identity (e.g., OPC, astrocyte markers), migration, vascular remodeling, and other glioma-associated traits. These changes suggest a shift away from plastic, invasive states toward a lineage-committed neuronal phenotype (Figure 4). The decreased expression of glial lineage markers supports the specificity and completeness of the reprogramming process.

#### 5. Role of Bioelectric Networks and Ion Channel Modulators in Neuronal Reprogramming

These networks, which spatially pattern resting membrane potentials during morphogenesis [61], are disrupted in cancer ecosystems, leading to depolarized cellular states and a loss of coordinated tissue organization. By targeting bioelectric networks with ion channel-modulating drugs, glioblastoma cells can be reprogrammed reducing their depolarized state and phenotypic plasticity [62]. The FID and FTT-treated samples exhibited strong upregulation of ion channel-related genes like GRIN1, GRIK1, NRXN3, and NRXN1, and calcium dynamics (e.g., CALB2), which regulate membrane potential and synaptic activity. In evidence, Liao et al. [24] recently demonstrated that glioblastoma cells can be transdifferentiated to neuron-like cell fates by upregulating NRXN1, one of the upregulated DEGs identified in our study herein. Further, they demonstrated that cell fate reprogramming significantly reduced astrocytic and mesenchymal-like states-driven tumor invasion and aggressive behaviors [24]. Thus, our findings highlight novel plasticity biomarkers and potential therapeutic targets for glioma cell fate reprogramming. Additional network analyses and heatmap visualizations (Figures S7–S8, Table S5) revealed bioelectric and synaptic regulators—particularly voltage-gated ion channels and neurodevelopmental genes—as key central nodes driving the neuronal differentiation.

### 6. Drug Targets and Precision Delivery Strategies for Cell Fate Reprogramming

This section expands on the molecular mechanisms and potential druggable targets referenced in the main discussion, focusing on the specific components and delivery strategies relevant to neuronal reprogramming therapies in glioma. For instance, the upregulation of WNT7A validates the specificity of the drug combinations used for neuronal reprogramming and underscores its role as a candidate target. Furthermore, ERBB3/4, a specific subtype of EGFR associated with pHGG aggressivity [11, 63-64], emerges as a precision therapy candidate in targeting glioma invasiveness. Moreover, a recent longitudinal single-cell analysis of pediatric high-grade gliomas identified ERBB2 and NRXN3 as key components of tumor–immune ligand-receptor interactions, including roles in synaptic signaling and growth modulation (Sussman et al., 2024). While that study focused on hemispheric pHGGs, our identification of ERBB3, ERBB4, and NRXN1/3 in brainstem H3K27M DMGs highlights a broader pattern of synaptic and neurodevelopmental reprogramming across spatially distinct glioma subtypes, supporting the hypothesis of convergent neurodevelopmental programs and stalled differentiation.
